# The separation of auditory experience and the temporal structure of MEG recorded brain activity

**DOI:** 10.1101/189480

**Authors:** Chris Allen

## Abstract

Do brain oscillations limit the temporal dynamics of experience? This pre-registered study used the separation of auditory stimuli to track perceptual experience and related this to oscillatory activity using magnetoencephalography. The rates at which auditory stimuli could be individuated matched the rates of oscillatory brain activity. Stimuli also entrained brain activity at the frequencies at which they were presented and a progression of high frequency gamma band events appeared to predict successful separation. These findings support a generalised function for brain oscillations, across frequency bands, in the alignment of activity to delineate representations.

## Main Text

The temporal structure of lived experience appears to be limited, where the minimum duration of individuated thoughts is approximately 100ms *(1, 2)*. This periodicity is coincident with the dominant brain oscillatory rate in the **a** range (~10Hz). Evidence from the visual and somatosensory domains, suggests that a function of specific frequency bands of oscillatory activity may be to individuate content (e.g. *3-5)*. However, there is a degree of inconsistency in the frequency specificity of these bands *(6)*. While this divergence may reflect task and modality specificities it is also possible that perceptual individuation exploits a more general and fundamental function of brain oscillations. Individuation of content requires representation over time. Brain oscillations may serve to facilitate representations by providing conditions where content can be delineated. If brain oscillations provide conditions for representation then, we would expect brain oscillations to place a limit upon how quickly one moment can be separated from the next according to innate oscillatory rate, thus constraining the flow of experience *(7-9)*. This possibility motivated a series of three pre-registered <u>(https://osf.io/h3z5n/)</u> hypotheses which related oscillatory brain activity measured using Magnetoencephalography (MEG) to how quickly one auditory perceptual object can be separated from the next.

The first hypothesis concerned the relationship between the prevalence of brain oscillations over a range of frequencies and capacity to individuate representations. The task involved presenting participants (n=20) with trains of noise bursts at different frequencies. The task was to count the number of bursts (between 4-7 bursts, see fig.1A,

https://osf.io/jz54p/). The stimuli were designed to merge when separation failed, resulting in lower performance. Therefore the stimuli probed capacity to separate auditory percepts over a range of temporal separations or frequencies. If a general function of brain oscillations is to provide conditions for representation over time, then completion of an oscillatory cycle may be required for such content to be separated. This meant that a relationship between the presence of oscillatory activity and task performance, over a range of frequencies, was predicted *(4, 10)*. That is, it should be possible to perform the task of counting bursts when there is sufficient time for an oscillatory cycle to have occurred and therefore burst separation performance should track the presence of oscillatory activity over a range of frequencies.

**Figure 1.**
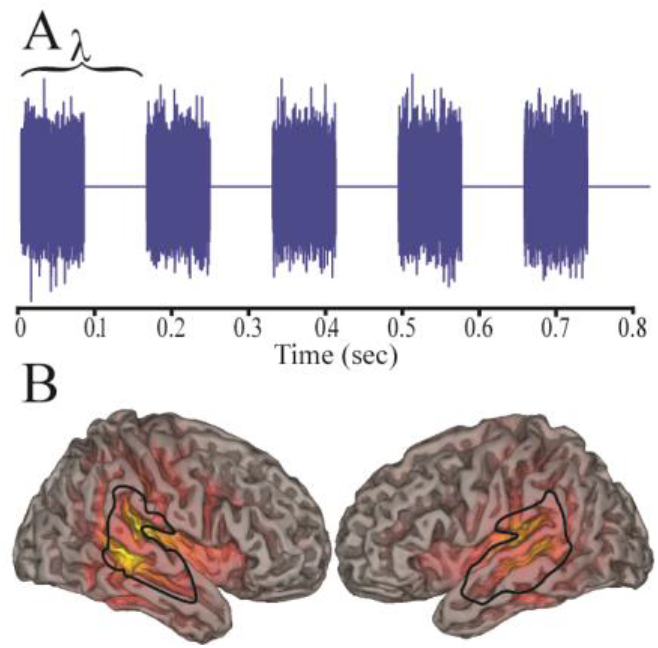
A. Illustration of an example burst train stimuli of five bursts. ***X*** refers to burst cycle duration, at 6Hz here. B. Representation of spatial distribution of group evoked response where brighter yellow colours indicate increased magnitude and the black line surrounds the location of the auditory cortex from which data was drawn (see methods).

The distribution of oscillatory brain activity was closely matched by the distribution of capacity to separate and count the numbers of bursts. Figure 2 illustrates the correspondence, over a range of frequencies, between behavioural performance and the magnitude of oscillatory activity (amplitude of the Hilbert envelope based on activity drawn from the auditory cortex using synthetic aperture magnetometry *(11)*). Statistically this was described as a within-subject correlation or ANCOVA *(12)*, where a strong predictive relationship between behaviour and oscillatory amplitude was expressed (r=0.88, F(1,219)=717.63,p=4.79×10-^71^,BF=2.81×10^13^). A correlation of this nature is to be expected when both measures conform to a 1/frequency distribution. However, an additional aspect of the analysis was the prediction that, if an oscillatory cycle is required to individuate content, then the behavioural distribution should overlay the oscillatory distribution, as plotted in Figure 2. Statistically this consistency was summarised by the ANCOVA’s intercept approximating to zero (value=-0.005(± 0.021SE), T(19)=-0.22, p=0.83, BF=0.24 see methods) where support for the null (BF<1/3) illustrates the match between the measures. A supplementary aspect to the analyses was whether differences between participants’ individuation performance related to their specific brain oscillatory make up, explored via the participant factor of the ANCOVA (F(19,219)=3.02,p=4.76×10-5, BF=6.92). Although this last comparison is a post-hoc exploratory analysis, it does suggest that differences in participants’ oscillatory brain activity may predict their specific ability to perform the task.

**Figure 2.**
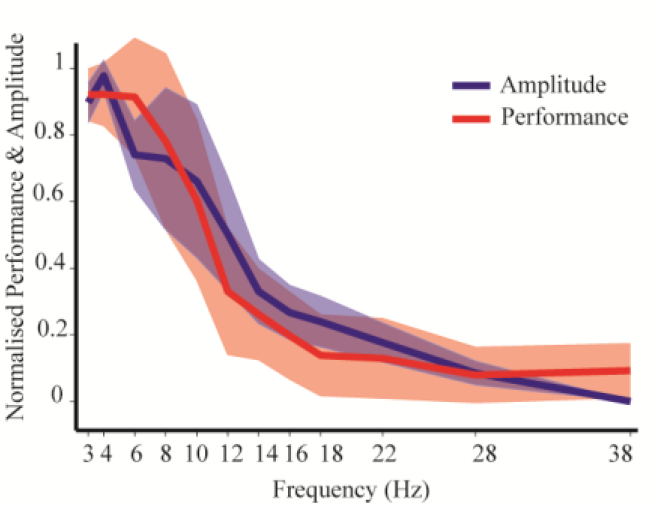
Group mean amplitude and behavioural performance against frequency (of presentation and oscillatory). Oscillatory amplitude was that of the Hilbert envelope applied to a virtual sensor activity trace constructed at participants’ peak evoked location within the auditory cortex. Performance refers to participants’ capacity to count the numbers of bursts at each frequency of presentation. Normalisation of both measures involved subtraction of minimum levels from their raw value and division by their range. Shaded areas represent 1 standard deviation.

The distributions of burst separation capacity and oscillatory activity bear a striking resemblance to one another, both conforming to what appears to be a 1/frequency (f) structure, prompting the question of the basis of this similarity. The amplitude 1/f can be understood as comprising the classic oscillatory frequency bands (a, (3 and y) and structured pink noise. The ubiquity and functional significance of scale free 1/f is controversial (13–15) and previous investigations of 1/f in brain and behaviour have often encompassed a much broader range of frequency distributions than those here (e.g.16). The current task sought to push perceptual systems to their limit in terms of how fast information could be reliably individuated, probing the lower limit of consecutive, separated, percepts. It found individuation closely matched rates of innate oscillatory activity. Therefore, although the behavioural and oscillatory data conform to what appears to be a 1/f distribution, the range over which they do so is limited, and is therefore not scale free. Instead, behavioural performance conforms to, and may be limited by, commonly observed oscillatory rates, which are, in turn, determined by the intrinsic size and conduction velocity properties of the brain (17). This relationship may be central to the similarity, correlation and intercept observed between the amplitude and behavioural performance distributions, suggesting that it is brain activity, most likely in the form of oscillations, which determine the rate at which percepts can be individuated.

The second main hypothesis and set of analyses concerned the effect the stimuli had upon oscillatory activity, asking if the stimuli entrain brain responses at the frequencies at which stimuli were presented (18, 19). The amplitude of the induced response was calculated for each frequency of presentation at the matching oscillatory frequency. These were then compared to the same amplitude measures, but with data drawn from trials where stimuli were presented at other non-matching frequencies, quantifying the brain’s frequency specific response to the stimuli. The pre-registered primary analysis comparing these conditions supported the prediction that stimuli entrained brain activity and this involved relative increases in oscillatory amplitude at the frequency at which the stimuli were (F(1,18)=10.22, p=0.005, BF=9.30, fig 3, see methods). Therefore the brain responded through entrainment to the stimuli frequency. This effect did not appear to be specific to any particular frequency range (validity × frequency interaction F(11,198)=1.05, p=0.40, BF=0.38, see methods).

**Figure 3.**
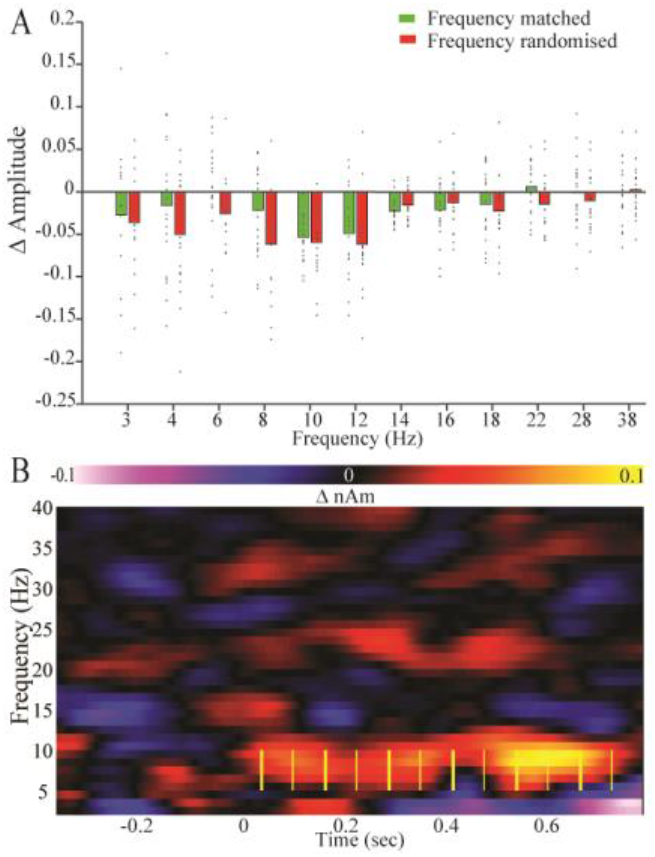
Entrainment analyses. A. Induced amplitude change, where data is drawn from either the matching frequency of presentation or where data was randomly down sampled from other frequencies. Bars are of group mean and data points are individual participants’ data. B. Example of group level time × frequency representations of differences described in A, yellow lines illustrate the stimuli (at 8Hz in this example, where thicker lines represent burst onset and thinner that of silent periods).

If oscillatory activity supports representation and therefore content individuation then we might expect a difference in oscillatory state between cases where individuation is successful vs. where it is not. The final set of analyses probed this hypothesis and consisted of three parts, testing different aspects of activity; oscillatory amplitude (induced and unbaselined), phase consistency and phase angle. Both oscillatory amplitude and phase angle expressed differences between data drawn from successful and unsuccessful trials in the y range. There was an amplitude desynchronisation in the successful condition predominantly prior to the onset of the stimuli (66-73Hz, from −356 to 18ms relative to stimulus onset, cluster p=0.018, fig.4B). This appeared to be followed, after the stimulus onset, by a relative synchronisation when pre-stimulus baselines were subtracted (66-73Hz, 7 to 678ms, cluster p=0.027). The angular phase difference preceded the amplitude desynchronisation from −626 to −313ms, at 66-77Hz. The statistical test applied here deviated from the pre-registered Wilson-Watson test owing to violations in assumptions. Instead Watson’s U2 tests were used, indicating an angular difference of ~ π, cluster p=0.037, fig.4A (20).

**Figure 4.**
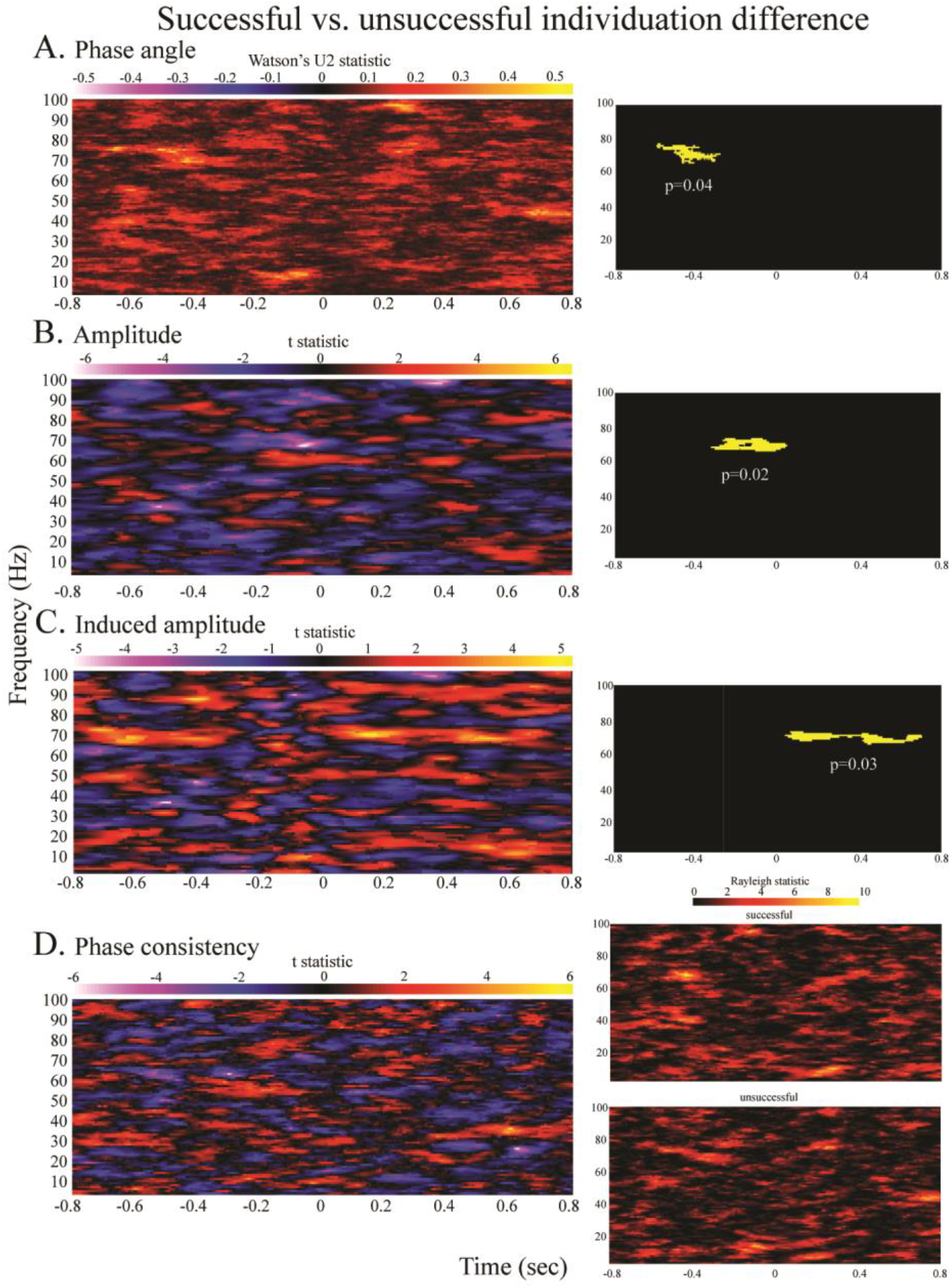
Time × frequency representations of group differences between successful and unsuccessful task performance in A) phase angle B) amplitude, C) amplitude where a pre-stimulus baseline was applied and D) phase consistency. Contrasts on the left are successful - unsuccessful. Significant clusters are indicated by yellow on black inserts. D represent the difference in phase consistency (phase locking value, left) and the levels of phase consistency in both conditions, represented by Rayleigh statistics (right).

This contrast between successful and unsuccessful burst separation highlighted a novel progression of narrow y band events starting with a phase difference, leading into a modulation in amplitude just prior to the stimuli onset, followed by a relative y response after the stimuli onset. These y events could be interpreted as being of approximately 9 duration, potentially reflecting cross frequency coupling (21). The pre-registered hypotheses predicted similar differences between the successful and unsuccessful conditions, with the observed sych/desynchronization directionalities, but these hypothesis also referred to differences within the a range based on previous research (e.g. 10, 22), therefore these findings are to some extent serendipitous. However, multiple studies have shown y band changes, particularly prior to demanding task elements (23, 24). The interpretation often applied is one of the allocation of top-down attention, expectancy or anticipation. Similarly it would appear that the state of preparedness is critical in determining whether or not bursts can be individuated and may be reflected by the y fluctuations. Speculatively, relatively fast y cycles may function to optimise the system’s susceptibility across a period, through increasing the opportunities for information transfer.

Together with the preceding analyses and previous demonstration of similar individuation dependent differences within a-p ranges (3-5), the y progression supports the interpretation that while band specific fluctuations may perform specific sub-functions these may also be subordinate to the more generalised function this experiment, and in particular the stimuli, were designed to reveal. The generalised function involves broad spectrum oscillations contributing to the delineation between the presence and absence of representations. For a neurone, or set of neurones within a population to exert influence and contribute to any particular representation, then just the right set of conditions must pertain, over time, which are conducive to that particular patterning of activity occurring (25, 26). Alternating through periods where suppression is imposed, followed by a relative relief from suppression brings ionic potentials into temporal alignment relative to threshold for action potential, thus imposing temporal structure, ordering and facilitating specific patterning and information transfer amongst and between populations (17, 27). This alignment can be interpreted as constraining potential outcomes (28), offering opportunities to bring neuronal assemblies into relief, against the background of neuronal noise. Transient formation of assemblies of this nature has been linked to the formation of conscious objects (9). Pushing the auditory perceptual system to its limit in terms of how quickly these assemblies or objects can be formed, dissolved and formed again was the aim of the stimuli design. The current findings relating the rate at which bursts could be separated and the rate of oscillatory activity therefore evidences the proposal that oscillations support to the delineation of representations and in doing so limit the flow of conscious experience.

## Methods

### Experimental Design

This experiment employed an auditory task in which participants were asked to count bursts of noise presented at different rates. This tested their capacity to individuate auditory percepts. At the same time oscillatory brain activity, localised to the auditory cortex, was quantified using magnetoencephalography (MEG). The theory under investigation was that brain oscillations may perform a function which allows for discrete individuated representation and therefore the separation of percepts. This led to a series of hypotheses concerning relationships between brain oscillations and the task which found support in the data.

The methods and materials for this experiment were registered prior to data collection and are available at: https://osf.io/h3z5n/. This section reflects the content of the online document as closely as possible. The methods described here encompass all procedures and analyses which were fully described and registered prior to data collection and any additions or alterations (beyond tense and clarification alterations) have been made explicit. A number of potential exploratory directions for analyses and theoretical considerations were noted and are still available via the original document. This article covers the primary registered analyses and hypothesis, but should readers wish to peruse the additional analyses, the data will be made available upon publication.

The materials and methods first describes the behavioural task in detail. This is followed by a description of the MEG acquisition parameters and the preliminary analysis methods. Primary pre-registered statistical analyses are then described. The exclusion criteria, examples of the participant instructions and post-task questionnaires are then provided.

All procedures were approved by the Research Ethics Committee at the School of Psychology, Cardiff University and all participants confirmed their informed consent to participate. Twenty participants completed the experiment, mean age 24.85 ±5.55SD, 14 female.

### The task

The behavioural task tested the segmentation of auditory information over a range of presentation frequencies. Trials involved exposing participants to between four and seven bursts of alternating white noise and silence. The periodicity of each burst and silence period was equal to the wavelength over a range of frequencies, from 3 to 38Hz (3,4,6,8,10,12,14,16,18,22,28,38 see fig. 1). 38Hz was chosen as the upper bound as it approaches the lower bound for perceiving the stimuli as an un-individuated tone (29). The white noise consisted of a random amplitude fluctuation at each sample time with a sample rate of 48000Hz (normally distributed, range (amplitude) adjusted according to participants’ preference, prior to the experiment). Participants were instructed to respond to the number of bursts of noise they heard within a train. This four alternative forced choice task measured participants’ ability to separate individual auditory percepts. Previous research relating audition to oscillations has tended to involve stimuli where the frequency structure may be ambiguous (e.g. ‘clicks’, where frequency could refer to the tone or the gaps in between or more complex sounds such as ‘chirps’ or speech (30–32)). Here, by contrast, the stimuli were designed to reflect a relatively simple dichotomous present/absent structure of oscillations over a range of frequencies. The idea which motivated this design was the possibility that oscillations may play a role in the delineation between the presence and absence of representations and that performing the task requires separation of representations.

Auditory stimuli were produced using Matlab (Mathworks Inc) in conjunction with the Psychophysics Toolbox and transmitted to the magnetically shielded room via E-A-Rtone (gold) insert earphones. Responses were collected via a Lumitouch (Photon Control) response box, where the index finger corresponded to four bursts, the middle finger to five, third finger to six and little finger to seven. See participant instructions below.

Participants completed four blocks of 144 trials during the experimental session, resulting in a total of 576 trials. Because some of the analyses required equal division of data according to behavioural success, as well as a possible division according to frequency of presentation, two of the central presentation rates were more prevalent than others to lend power to these analyses. Within each behavioural block all 12 frequencies were presented at least eight times. As there were between four and seven bursts at each frequency, each unique stimulus was presented at minimum twice per block. The frequencies presented more often were at 10 and 14 Hz as pilot data suggested that these approximate the median range of behavioural performance where success is approximately 50%. Each of these frequencies was presented 32 times per block. Trial orders were randomised within each block. Trials were separated by a 1.25 second interval following competition of the non-speeded response. Short breaks (~1-2 minutes) were given in between blocks to reduce fatigue, although subjects were asked to maintain head position throughout collection of experimental data. Prior to the collection of MEG data, subjects were familiarised with the task through the completion of 1-5 behavioural blocks of varying length. To reduce scanner time this familiarisation was carried out using different equipment and often during a separate session. No data was retained from the familiarisation.

### Data acquisition and preliminary data analyses

This experiment used many of the same participants and collected data during the same session as another pre-registered MEG experiment. The current experiment was performed after a stop signal task, the details of which can be found at https://osf.io/r8tvu/. Participants were given a lengthy break between experiments.

MEG data was collected on a 275 channel CTF system, sampling at 1200Hz. Data was collected in the supine position. Participants were instructed to maintain the same head position, to the best of their ability, following the initial head localisation procedure, used for all subsequent blocks.

For the majority of analyses and source localisation steps, data was epoched into individual trials in sections from −800 to 800ms relative to the onset of the digital trigger for the first stimulus. This meant that a later portion of the full burst trains was not captured for some of the lower frequencies of presentation, when trains contained higher numbers of bursts. The reason for this was to allow for consistent temporal structure of trials without introducing long intervals between trials at higher frequencies with lower numbers of bursts, as this had the potential to impact upon performance and data quality. This individuated all trials without the inclusion of multiple trials into a single epoch and should not have biased comparisons across frequencies of presentation. Individuated trials were visually inspected, with the experimenter blind to frequency of presentation and success status, and clearly corrupted trials (e.g. by movement) were excluded for the analysis. This resulted in the inclusion of 10739 trials out of a possible 11520.

Pilot data and previous research (e.g. 33) suggested that brain responses to the onset of the first burst of trials have sources that approximate to the auditory cortex when analysed using whole brain Synthetic Aperture Magnetometry (SAM 34). The evoked change, independent of non-phase locked oscillatory changes, was taken to be representative of the cortical auditory response and used to construct an auditory ‘virtual sensor’ (34). This was then used to analyse the oscillatory activity in subsequent analyses. Although evoked changes could potentially be independent of oscillatory changes of interest here, this method should have targeted biologically plausible cortical auditory responses, and more importantly, should not have biased the analysis to a particular frequency range (for example, if induced oscillatory SAMs were used, rather than evoked SAMs, the location of the virtual sensor would likely have been driven by a oscillations due to 1/frequency distribution). The spatial location of this virtual sensor was further constrained to the auditory cortex using a binary mask derived from the Harvard-Oxford atlas (35) representation of the superior temporal gyrus (composed of both anterior and posterior sections), thresholded to 25% (e.g. 36). The peak activation within these masks, during a 50 to 400ms active period following the onset of the auditory stimulus, over all trials, was isolated for each participant using mri3dX (Singh KD) and used to produce activity traces. The baseline applied to produce the evoked data comprised the final 350ms prior to the onset of the auditory burst trigger and the evoked SAM made use of a 200Hz low pass filter. This filter was broader than is commonly used, again to avoid biasing the sources to any particular oscillatory frequency response. All filters are zero phase bi-directional. A 1Hz high pass filter was also registered, but this was never applied as it could have suppressed evoked responses of interest. Also, in addition to registered methods, all analyses using baseline or active periods applied a +30ms offset to timings to accommodate for the transmission from the E-A-Rtone modals to ear inserts (~1m).

Spectral oscillatory amplitude and phase information was estimated using a Hilbert transform applied to the resultant virtual sensor between 3 and 100Hz with a step size of 1Hz and bandwidth of 6Hz. Unless otherwise stated, the same temporal parameters as described above for the evoked SAM were used to produce time × frequency spectra upon which subsequent analyses were based.

### Statistical approach

There were three pre-registered analyses applied to this data which are described in this section. The first assessed the correspondence between the spectral MEG data and behavioural ability. The second probed the nature of the brain response to individual bursts of noise at different frequencies of presentation. The final analysis method tested differences in cortical responses between when subjects were able to segment information and when they failed to do so and asked if such differences were predictive of success. An additional exploration of the topography of the observed differences is then described. Participants also completed post-task questionnaires, responses to which were related to the MEG data in a series of exploratory analyses, which were developed after data collection and are described at the end of the materials and methods.

### Correlational test between oscillatory cortical response and behaviour

The auditory bursts were designed to merge if efficient segmentation was not possible, thereby imposing a limitation upon behavioural performance which should track the neuronal mechanisms of content individuation. If brain oscillations act to individuate content, then the expectation was to observe a direct correspondence between the range of frequencies over which brain oscillations occurred and the rate (frequency) at which participants were able to individuate content and perform the task. Therefore, if cortical oscillatory activity fulfils the function of cognitive segmentation, then the expectation was that there may be a direct correlation between oscillatory activity and behavioural performance over a range of frequencies.

A central aim of this analysis was to explore the correspondence between behavioural performance in the auditory task and the MEG data. The amplitude of the Hilbert envelope was representative of the presence of cortical oscillatory activity at each frequency of interest in the MEG data. As a point of clarification, the term ‘oscillatory’ is linked to the amplitude of the Hilbert envelope here. However, it has been suggested that such a measure also represents structured pink noise (37). This could bring into question the interpretation of the primary correlation being between oscillations and behaviour. That said, the structured pink noise still denotes underling activity with periodicity that repeats at the frequencies of central interest to this investigation and correspond to the behavioural distribution, which can be subsumed under a broad interpretation of the term oscillatory.

Through a series of correlations, the first set of analyses therefore tested the relationship between task performance and oscillatory amplitude localised to the auditory cortex. The behavioural data was the normalised proportion correct with data points at each frequency of presentation. At each of these data points the absolute amplitude of the Hilbert envelope at the corresponding cortical frequency was the primary MEG dependent measure, which was also normalised to 0-1. To clarify, both behavioural and amplitude measures were normalised for each participant by subtraction of the minimum value and division by their range. The derivation of the oscillatory amplitude used all non-excluded trials and applied the same parameters in terms of active periods as used in the construction of the evoked trace, where the mean amplitude was taken across the active period (50 to 400ms). Also, in clarification of the pre-registration document; as this analysis concerns a quantification of the relative presence of oscillations at different frequencies, no pre-stimulus baseline was applied to the main analysis. However, as the oscillatory makeup in response to stimuli may also be of interest, the same analyses as described here were repeated to explore the induced response where pre-stimulus baseline was subtracted, the results of which are described below in supplementary text.

The first correlation simply asked the extent to which the behavioural data was predictive of MEG amplitude data and vice versa. This involved matching behavioural performance over the range of frequencies of presentation to the normalised amplitude of the Hilbert envelope over the same range of frequencies, and applying an analysis of covariance to access the correspondence between the two, while treating individual subjects as partialled out categorical variables (12). To clarify, the corresponding Bayes factor (BF) was the result of applying the JZS prior with default scaling (0.7071) to the t-statistic of the regression slop (38).

The resulting correlation coefficients were then used to characterise relationships between the oscillatory and behavioural data in secondary analyses. If the stimuli allow us to track content individuation, and quantifiable oscillatory activity performs its proposed role (i.e. stimuli can only be individuated if a full oscillatory cycle has occurred), then it was expected that the intercept of the correlations would be roughly zero. Conversely if more than one oscillatory cycle is required to individuate a percept, then the expectation was that the intercept coefficient, on the behavioural axis, should be positive. If less than an oscillatory cycle was required, the intercept coefficient should be negative. These possibilities were tested with a Bayesian equivalent to a T-Test (38) where coefficients consistently around zero would have been expressed by BF<1/3, a consistent fluctuation in either direction as expressed by BF>3, and inconclusive data would have been expressed by BF~1. An additional ANOVA of residual error was described in the pre-registration document. This was aimed at exploration of any discrepancy between the measures over the frequencies under investigation. However, as the primary analysis of the intercept supported the absence of their bringing a discrepancy, the corresponding ANOVA was redundant and therefore not applied.

The primary set of tests can, therefore, be interpreted as a set of within-subject correlations or analyses of covariance (ANCOVA). Also, of secondary interest here were the between-subject differences, i.e. whether differences in individual participants’ performance of the task corresponded to differences in their oscillatory brain activity. This question was posed by collapsing the behavioural and spectral data across frequencies and testing the correlation between the two means, across participants *(39)*, using a Pearson’s correlation and complimentary Bayesian correlation *(40)*. This question was also posed at each frequency of interest (see table ST1). Resultant r values were compared to zero with a single sampled t-test (and equivalent Bayesian test), and together with an ANOVA applied across frequencies, had the potential to reveal frequency specificities of individual difference relationships between measures. The results of these tests are presented in supplementary text below. Additionally it became apparent, after study registration, that the variability in the relationship between the behavioural and oscillatory data which pertained to individual differences was summarised more succinctly, directly and utilising more of the data, in the participant factor of the primary ANCOVA, than has been reported in the main text.

Following registration of this experiment and data collection it also became apparent that the 1/frequency distribution observed in both the behavioural and oscillatory data can be summarised by a single coefficient **p** within the formulation: Amplitude or Behaviour ≈1/frequency*(41, 42)*. 1/ frequency curves were therefore fitted to the group mean oscillatory and behavioural data, using Matlab’s curve fitting toolbox with default randomised initial fitting parameters. The similarity or difference between oscillatory and behavioural **p** coefficient were then assessed with a paired t-test and Bayesian equivalent *(38)*. Additionally, as an exploratory analysis based on previous research *(42)*, individual participants’ behavioural and oscillatory **p** coefficients were correlated with one another.

The prediction of this section, based on the idea that innate brain oscillations determine the minimum duration of the individuation of content, was that there may be a direct correspondence between the distribution of oscillatory amplitude over a range of frequencies, and behavioural performance over the range of frequencies of presentation.

### The effect of the stimuli upon oscillatory cortical response

The second series of analyses posed the question of whether or not oscillatory amplitude is modulated and entrained by the frequency of presentation (e.g. *43, 19)*. As this question pertained to the frequency at which stimuli were presented, the data was divided accordingly into frequency specific data sets. The effect of presentation frequency was assessed by deriving point values for oscillatory amplitude, within each frequency specific data set, at the corresponding frequency of presentation. These were then compared to point values based upon the induced response at the same oscillatory frequency, but where MEG data was drawn from all other frequencies of presentation. For example, where stimuli were presented at 8Hz the point value was the mean amplitude at 8Hz across the active period, and this was compared to the amplitude at 8Hz but where the contributing data was from trials where the stimuli had been presented at 3Hz, 4Hz, 6Hz, 10Hz etc. Data sets used in this contrast were randomly downsampled so that trial numbers were equal across the comparison (see below).

The question here relates to brain responses across the course of the train of stimuli. Therefore the temporal structure of the data differed from that described in the data acquisition section above. The active period was from the onset of the first burst until 200ms after the end of the final burst within a train. The baseline period covered an equal period, prior to stimuli onset, offset by 50ms (see fig. S1). These periods were also limited to contain no more than 800ms of data, so as to avoid inclusion of previous trial data.

Whether induced responses are modulated by frequency of stimuli presentation, and if such a relationship holds over a range of frequencies, was tested with a repeated measures ANOVA. The dependent variables were the point values for oscillatory amplitudes. The factors for the ANOVA were whether or not data was drawn from the corresponding frequency of presentation (termed ‘validity’, 2 levels) and the frequency of presentation (12 levels). A significant main effect of validity probed the primary question of this section and indicated that the frequency of presentation had a direct effect upon the corresponding oscillatory brain response. The application of Chauvenet’s criterion *(44)* was registered, as clarification it was applied to difference tests with a default Criterion level of 0.5, and here it was applied to the main validity comparison resulting in the exclusion of one participant’s data form the main entrainment analysis. A statistical interaction involving the frequency of presentation factor would have suggested that the effect may have been present over some frequencies and not others. Bayes factors reported are the result of the integration of JZS priors with default scaling using the relevant F and T statistics *(38, 45)*.

Following registration a discrepancy of interpretation between frequentist and Bayesian ANOVA and t-tests was noted which requires clarification. Under a frequentist approach the simple main effect of a two level factor (e.g. validity) of an ANOVA is equivalent to a t-test applied to the means across other conditions, where F=√T. This is not the case for default Bayesian ANOVA’s *(46)*, where differences within, or the presence of, factors that are not of interest to the primary question, effects the outcome of the primary simple main effect analysis. This makes interpretation difficult. Therefore the BFs reported in the main text are the result of applying the Bayesian T-test to mean differences between conditions where there is no inconsistency in interpretation *(38, 45)* and these are supplemented for completeness by the results of the Bayesian ANOVA *(46)* below. The results of the two approaches were broadly consistent and differences concerned the relative weight of evidence.

In the interest of transparency the following additions to the registered procedure are noted. Initial inspection of the data indicated a significant effect of validity (F_(1,19)_=12.88, p=0.002, BF=20.45), however the temporal parameters used in this computation were incorrect by 30ms (the time allowed for stimuli to reach participants’ ears from the speakers had been omitted). Therefore it was necessary to recompute the analysis, the results of which are reported in the main text. In doing so it became apparent that the result was highly dependent on the registered randomised downsampling reapplied in the construction of the invalid contrast data set. To avoid this issue an additional analysis was undertaken where all trials were used, the results of which are reported in the supplementary text as explicitly exploratory and are not susceptible to randomisation effects. The resultant contrast resulted in 536.95 ± SD47.23 mean number of trials for each participant in the valid condition and 5906.45 ±SD 519.57 in the invalid condition. This compares to the mean of 536.95 ± SD47.23 trials in both conditions under the pre-registered downsampling approach.

The prediction of this section was that if oscillatory cycles fulfilled the proposed role of functional segmentation, then the expectation was that brain oscillations would be modulated by frequency of stimuli presentation in such a way that they would shift to match the frequency of presentation, allowing segmentation. Therefore the prediction was that the amplitude of induced response would be increased at the frequency of presentation.

**Figure S1.**
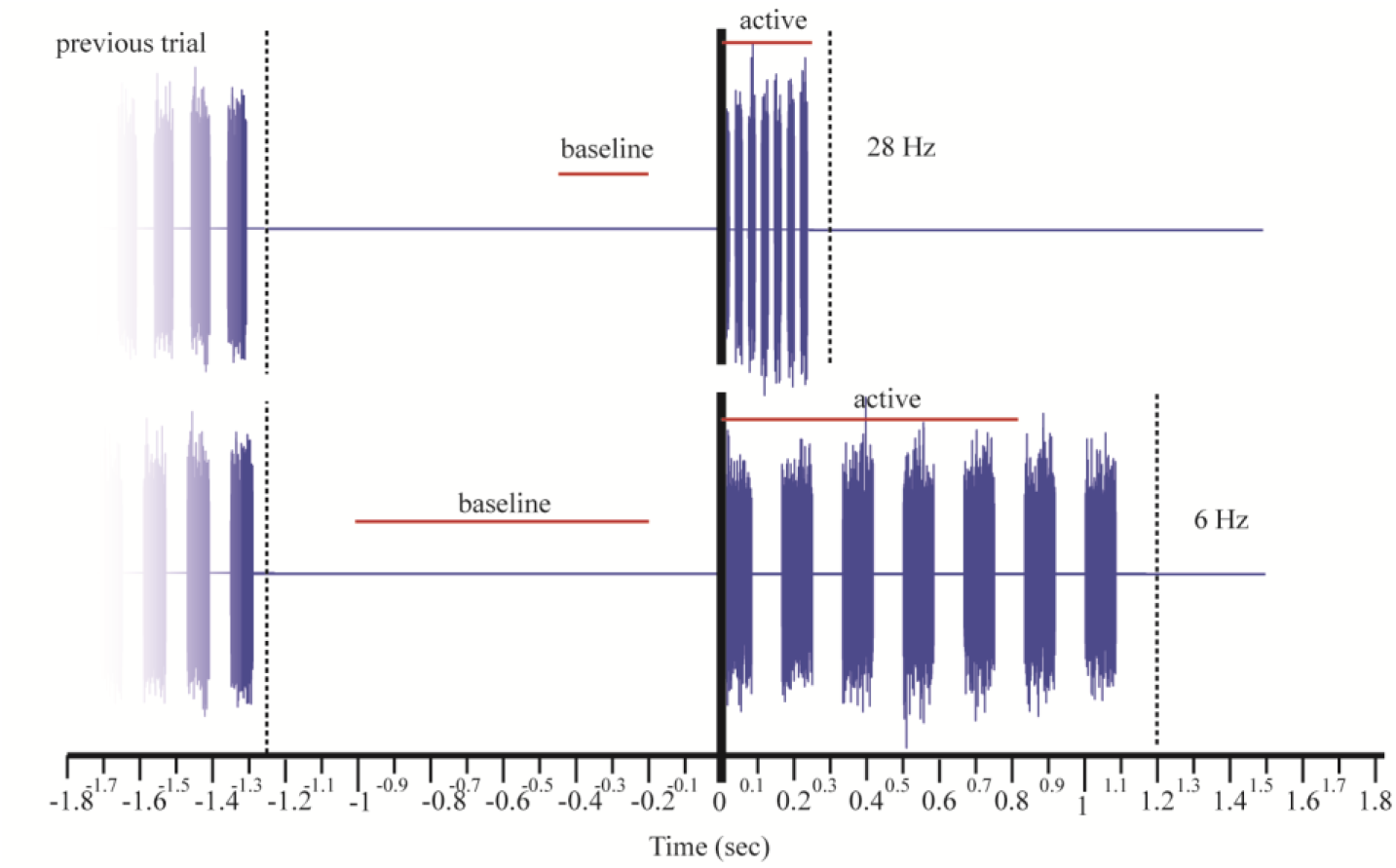
Illustration of temporal division of data applied in analysis of the effect of the stimuli upon oscillatory cortical response. Two trials are depicted where stimuli are presented at 28 and 6 Hz. Dotted lines indicate separate trials.

### Are changes in oscillatory cortical response predictive of behavioural success?

If brain oscillations fulfil their proposed role in segmentation, not only should the frequency of presentation influence the brain’s oscillatory response (above) but the oscillatory state may also predict the success of behavioural performance. Of central interest here was the relationship between behavioural success and cortical response. The analyses described in this section correlated the extent to which both oscillatory amplitude and phase were predictive of behavioural success. This contains three interrelated subsections which address different aspects of oscillatory activity, relative to behavioural success. The first concerns oscillatory amplitude, the second tested phase constancy and the third analysed phase angle.

These analyses were applied to the higher prevalence trials (10 and 14 Hz). Two data sets were derived for each subject, one comprising successfully segmented trials and the other unsuccessfully segmented trials. Randomised downsampling was applied once so that for each participant an equal number of trials contributed to each data set. In accordance with pre-registered criteria the high prevalence data sets were merged to meet the requirement of a minimum of 20 trials per condition. This meant that it was not possible to assess the interaction between behavioural success and frequency of presentation within this data. The temporal structure of the data followed the description in the statistical data acquisition section. Temporal baselines were not applied for these analyses, with the exception of the analysis of oscillatory amplitude which was tested with and without the baseline described in the data acquisition section.

### Oscillatory amplitude

Quantification of the predictive power of these data sets with respect to behavioural success was applied through a series of permutation based tests upon the oscillatory amplitude. These ask, in both the time and frequency domain, where a consistent difference between successful and unsuccessful conditions might be found. Oscillatory amplitude was quantified using the Hilbert transform as described in the data acquisition section. The question posed here was applied to each frequency and time point within a range from −800ms to 800ms relative to the onset of the first burst trigger. The frequency ranges used was 3:100Hz (1Hz step size). The secondary exploration of a more restricted frequency set (3:18Hz) described in the original pre-registration document was unnecessary owing to the differences apparent within the primary analysis.

Primary analyses here were posed at the group level, using the variance across participants. The contrast between successfully segmented trials and unsuccessful trials was conducted with a series of permutation tests where the adjustment for multiple comparisons applied a mass cluster-based approach *(47, 48)*. The cluster-based correction followed the principles described by *(49)* and used the maximum cluster mass method based on the sum of the t-statistics with Chauvenet’s correction. 5000 permutations were used and 8-way connectivity over time and frequency was used to isolate clustered differences (criteria for inclusion: p<0.05). Supplementary to the cluster-based correction, a false discovery rate *(50, 22)* correction was described in the pre-registration document and was applied to the p values derived from the time × frequency comparisons described. However, there are violations of the independence assumption of FDR correction, when applied to the time × frequency data. The option of an amendment to overcome this dependency *(51)* did not seem appropriate as this would have penalised analyses for adjoined, correlated expressions of differences, which are theoretically more plausible than separate, independent differences. The violation of independence does however mean an emphasis should be placed on the supplementary and exploratory nature of the FDR-corrected tests, reported below in supplementary text.

Individual participants’ data were also analysed in a secondary set of analyses where variance was drawn across trials, rather than participants (fig. S7-11, table ST2), the results of which are presented in supplementary text below. At the individual participant level a deviation from the pre-registered protocol was applied, reducing the numbers of permutations to 1000. This was due to feasibility as the computational demands, of the phase analyses in particular, are high (relying upon permutation tests imbedded within permutation tests, see below). The effect of this reduction, if anything, is to make these analyses more conservative. A supplementary point value analysis described in the pre-registration document was not applicable here owing to the merging of high prevalence data sets, which meant that there was no specific frequency from which to draw point values. This, in turn, was necessitated by the registered criteria which required a minimum of 20 trials in each condition.

The predictions made with respect to this section were that if an oscillatory cycle is required to successfully segment information then, because an oscillation will have occurred, oscillatory amplitude should be higher on successful trials compared with unsuccessful trials. An additional expectation was that if oscillatory cycles were required to perform the task successfully, then the absence of such cycles prior to the presentation of stimuli might be conducive to successful performance. Therefore a relative desynchronisation of oscillatory responses prior to or during stimuli onset, in the successful condition, was predicted as described by *(22, 52)* and that such an effect might have been prevalent in the α range.

### Phase consistency

The question posed by this series of analyses was whether there was consistency in the phase angle of oscillations when trials were successfully segmented, when they were not, and if there was any difference in consistency between these two conditions. Rayleigh tests were applied to successful and unsuccessful data sets separately *(20)*. At the group level these used the circular mean phase angle across trials *(20)*. These assessed if the data expressed phase consistency (H1) or were uniformly distributed (H0). Then the difference in consistency was accessed through the comparison of a quantification of phase-locking (known as the phase-locking value, PLV see Lachaux et al, *(53)* for details, note here the PLV was computed across trials rather than sites and the Hilbert transform was used, as opposed to a complex Gabor wavelet), which can be described as follows:

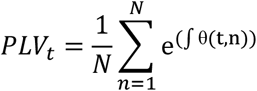

where t are the time points, n are the trials or group (n[1,…,N]) and θ(t, n) is the angular difference across sets of trials (ø_1_**-**….ø_n_). Hence perfectly aligned phase angles across trials will have a PLV value of 1 and approach 0 as phase consistency is lost. The same analysis method as described for oscillatory amplitude was then applied here to assess if there was a difference between conditions. That is, a set of difference scores (of the PLV here) were computed based on the differences in data drawn from successful and unsuccessful data sets. PLV comparisons were applied using the t-test related cluster statistics previously described.

At the participant level, unlike the other measures, it was not possible to compare the phase consistency measures as the measures used the variance across trials in their construction. Also it was not possible to apply the mass cluster-based correlation for multiple comparisons to the Rayleigh tests as these tests did not involve a direct comparison between successful and unsuccessful conditions. These two points are clarifications and additional to the original pre-registered document.

The hypotheses here were that phase may be more consistent for successful trials than unsuccessful, resulting in a positive PLV difference, and in the time × frequency domain this may be observed as a predictor of success shortly before the onset of the stimuli, in the lower frequency range (based upon *(22))*.

### Phase angle

These analyses tested the difference in angular phase of oscillations in both the successful and unsuccessful conditions. As this question related to angular distribution, a series of Watson-Williams tests *(20)* were applied in each of the individual participant’s data sets, using the variance across trials. At the group level, using the circular mean phase angles across trials *(20)*, drawing variance across the group of participants, the same cluster-based permutation tests were applied to see if any differences in mean phase angle were expressed. These would have effectively applied a Watson-Williams test at each time and frequency of interest (parameters as described above) and, using the p statistic, a cluster-wise and FDR correction. However, following data collection, it was found that in many instances the assumption of homogeneity was violated *(54)* and therefore the application of Watson-Williams tests was unviable. Following this, the Watson-Williams test was replaced by the non-parametric Watson’s U2 test, as recommended by Zar *(20)*. This involved permutation testing at each time and frequency point where, in keeping with the cluster-based permutation testing, 5000 permutations were used at the group level and 1000 at the participant level. The statistic used for the mass cluster calculation was therefore the Watson’s U2 statistic.

The prediction here was that behavioural success may have been more prevalent when the onset of stimuli were coincident with the negative trough of oscillatory cycles within the lower frequency ranges *(55, 4)*.

It is possible that similar randomisation effects as those reported with respect to the entrainment analysis could have affected these analyses (see *the effect of the stimuli upon oscillatory cortical response* section). However, the use of all trials here, as used in the entrainment analysis to alleviate this concern, would have caused substantive violations to the assumptions of the majority registered analyses of this section, particularly those that involved permutations of trial conditions (i.e. Watson’s U2 static and mass cluster based corrections) and those involving quantification of phase locking. Of the tests comparing successful and unsuccessful trials, this left only the amplitude and PLV t-static based difference, where FDR correction for multiple comparisons was applied, as the set of tests which could be applied and utilised all trials. These tests between all successful vs. unsuccessful trails where therefore applied and are reported in the supplementary text as additional exploratory analyses.

### Exploration of the topography of behavioural success differences

Following the collection of the data the analyses revealed a series of **y** band differences between successful and unsuccessful burst counting, a potential concern being that these could pertain to muscle artefacts. Although obvious muscle artefacts had been removed from the data a further set of exploratory analyses were undertaken to assess the spatial distribution of the differences *(56)*. This applied the SAM technique *(34)* described above in the oscillatory domain where the parameters corresponded to the **y** differences established using the pre-registered methods. The pre-stimulus **y** desynchronisation covered a period from −356 to −18ms between 66 and 73Hz. The SAM analysis applied to the induced difference was baselined (-350 to 0ms), again used a 66-73Hz frequency range and an active period from 7 to 678ms relative to stimulus onset. At present I am not aware of comparable spatial techniques having been developed for angular phase differences, so the topography of **y** phase differences were not assessed in a corresponding way.

### Exclusion criteria

Participants were to be excluded whose data indicated excessive movement during acquisition. This may be detected by a greater than 10mm difference in head position between acquisition blocks or more than 10% of trials being excluded on the basis of muscle/movement artefacts within any given block, as detected during initial data inspection.
Participants who reported excessive fatigue or discomfort during acquisition were to be excluded, as would participants who fell asleep during acquisition.
Behavioural data that indicated participants did not perform the task as instructed (e.g. fingers allocated to the wrong response buttons) would have led to the exclusion of the corresponding data.
Other unanticipated technical failures may have led to the exclusion of data.
Participants were free to withdraw their participation and data at any point of the investigation.

One participant was excluded on the basis of excessive head movement and self-reported excessive fatigue. The corresponding data was not analysed nor was it included in the 20 participants described above.

### Participant instructions

The following instructions were given to participants in written form before starting the experiment.

You will be presented with bursts of noise, in trains of between 4 and 7 bursts. These burst trains will be at different rates. Your task is to *count* the number of bursts.

The response box has 4 buttons on its top which correspond to responses 4, 5, 6 and 7 going from left to right. You can take your time to respond and the next train won’t start until shortly after you have made your response to the previous train of bursts. There are to be 4 blocks of burst trains each lasting approximately 8 minutes.

When doing this task please close your eyes, try to stay awake, try not to move and count the number of bursts.

### Post task monitoring

Participants were asked to complete a questionnaire after the MEG data collection. The analyses applied to the questions related responses to the MEG data through a series of correlations. MEG variables were the primary measures of the three main sections where differences in the MEG data were apparent. This meant that these analyses were developed after the collection of data. Therefore, although the use of the questionnaires was pre-registered, all related analyses were developed post-hoc, and as such, are to be considered as exploratory.

The measures drawn from the main three analyses sections are as follows: The first main analyses probed the match between the behavioural distribution and the oscillatory amplitude distribution. The estimates of the extent to which individual participants displayed a correspondence between the behavioural and oscillatory distributions are summarised by their individual r coefficients and gradient (m) values. The offset between the behavioural and oscillatory distributions is summarised by the intercept (c). Correlations were also applied between the questionnaire responses and the β coefficients from the 1/frequency^p^ fits derived for behavioural or oscillatory measures separately, as well as the discrepancy between the behavioural or oscillatory distributions summarised in a β difference score. The second main analysis probed the entrainment effect which can be summarised for each participant as a single measure of the difference in amplitude between where data is drawn from the matching/valid frequency and the downsampled randomised alternative set of trials. The final set of main analyses contrasted successfully individuated trials against unsuccessful trials and showed appreciable differences at the group level in three domains: raw amplitude, induced amplitude and phase angle. As the phase data is circular the correlation was restricted to amplitude differences. This used the mean amplitude difference between the successful and unsuccessful conditions across the time and frequency regions highlighted by the mass cluster-based correction (depicted in fig. 4).

Prior to analysing the relationships between the questionnaire and MEG data the following hypotheses (HQ) were drawn, for each of the questions posed to the participants. The questions (Q) as presented to the participants are given below in italics. These are presented together with the corresponding hypotheses. Coding of responses has also been provided. Numbering of questions, hypothesis and the coding have been added here for clarification but were not presented to participants. Correlations between questionnaire response and measures described above used Pearson correlation for continuous responses and Spearman’s for categorical responses, the resultant r values were then used to apply a Bayesian correlation test *(40)*.

Q1. 1A. *Did you find yourself counting the number of bursts in your head?*

*Yes [ ] No [ ]* (yes=1, no=0)

1B. *If so please make a mark on the line below to estimate how often that you adopted this strategy?*

*0% 50% 100%*

*None of the time Half of the time All of the time*

(as proportion 0-1)

HQ1. The task was designed to rely upon such a strategy of counting to individuate bursts. Therefore participants who “counted the bursts in their head” (response 1 question 1A and higher responses to question 1B) may have expressed larger effects of the stimuli in terms of both the entrainment effect and differentiation between successful and unsuccessful trials. Furthermore, as the proposal is that task performance realise upon such a strategy and this may have been facilitated by brain oscillations, it was predicted that the coupling (r and m values, of individual subjects correlations of the primary ANCOVA) may be higher in such participants.

Q2. 2A. *Did you count as the bursts were presented or look back / reflect upon the stimuli afterwards?*

Count as they were presented [ ] Reflect afterwards [ ]

(reflected back=1, as presented=2, both=3)

2B. *If you reflected upon the trains afterwards, how often would you estimate that you adopted this strategy?*

*0% 50% 100%*

*None of the time Half of the time All of the time*

(as proportion 0-1)

HQ2. Participants who counted the bursts as they went along, as opposed to those who reflected back after, (response 1 for question 2A and lower responses to question 2B) might have expressed larger entrainment effects and a greater differentiation between successful and unsuccessful trials as both analyses were orientated towards differences around the period of stimuli presentation.

Q3. *3A. Did you ever find yourself estimating the number of bursts based on the duration of the train rather than count the bursts individually?*

*Yes [ ] No [ ]* (yes=1, no=0)

3B. *If so, how often would you estimate that you adopted this strategy?*

*0% 50% 100%*

*None of the time Half of the time All of the time*

(as proportion 0-1)

3C. *Also with respect to when you adopted this strategy, was it for;* all trials regardless of rate [ ], just fast trials [ ], just slow trials [ ].

(All=1, fast=2, slow=3)

HQ3. Participants who based their responses on the duration of the burst trains rather than individuated burst (question 3A response 1 and higher responses to question 3B) might be expected to show weaker entrainment effects and differences between successful and unsuccessful trials, again owing to the temporal structure of the analyses; being based around the period surrounding burst onset. Perhaps more importantly, participants who used the train duration might be expected to have expressed a reduced coupling between the oscillatory and behavioural response distributions assayed in the first analyses. The reason being that estimation of duration may be facilitated at higher frequencies, where duration is shorter *(57)*. Adoption of this approach may have resulted in reduced correlation (lower r values) and potentially negative gradients (m values) for the fits between behavioural and oscillatory data. This might practically be the case if participants were to perform the task based on duration of trains when they are presented at lower, or across, frequencies of presentation (responses 1 and 3 to question 3C).

Q4. 4. *Please make a mark on the line below to indicate what you did when you did not know how many bursts there were. Did you*

*Always give 50/50 Just press your best guess? mix any button?* (always best guess 0-1 any button)

HQ4. It is possible that participants who gave “their best guess” best guess when they were unsure as to the number of bursts, as opposed to “just pressing any button” (lower responses to question 4), may have utilised some form of residual or unconscious capacity (e.g. *32)*. This might have increased their capacity resulting in lower behavioural β coefficients when exponential models were fitted. If, however, the oscillations perform a role in specifically conscious processing, then we might expect the discrepancy between the oscillatory and behavioural data (c intercept of analysis 1) to be greater in these participants who may utilise unconscious capacity.

Q5. 5. *Please describe any other strategy you used, especially if the above questions do not provided an adequate description*.

Please give any other comments you wish to make.

No specific hypotheses or analyses were targeted at the final question.

## Supplementary Text

This section supplements the main text describing exploratory and registered analyses and alternative representations of the data. It follows the structure of the analyses in the pre-registration document and main text, consisting of three sections pertaining to: i) the relationship between oscillatory and behavioural distributions, ii) the entrainment of oscillations by stimuli and iii) the divisions of data according to behavioural success. These are then followed by a description of the results relating to the questionnaire data.

### Relationship between oscillatory and behavioural distributions

An alternative graphical representation of the primary ANCOVA relating the oscillatory and behavioural data is presented in fig. S2 and S3 *(12)*. These uses the same data as fig. 2 of the main text but provides a more complete, although less intuitively interpretable, representation of the data. An alternative exploratory statistic which summarises the correspondence between the oscillatory and behavioural data is the interclass correlation coefficient *(59)* applied to the ANCOVA which was 0.956. Raw behavioural performance levels as a proportion correct ranged between 0.92 (±0.11SD) at 4 Hz and 0.29(±0.08SD) at 28Hz where chance performance is 0.25.

The primary analyses of this section investigated the relationships between behavioural performance and the raw oscillatory amplitude quantified via the application of a Hilbert envelope. Additionally, the same main analyses were applied but where a pre-stimulus baseline was subtracted in order to test the induced oscillatory response to the onset of the stimuli. In this exploratory analysis the general partnering of the data, together with the strong correspondence between the oscillatory and behavioural distributions, was maintained, although the effect size was slightly reduced (r=0.56, F_(1,219)_=100.60,p=1.01 ×10^-19^,BF=2.38×10^6^). Reduced in effect magnitude can be expected following the subtraction of the baseline which itself contains random fluctuations. Interestingly the oscillatory distribution of this induced change from pre-stimulus levels was now to the left of the behavioural distribution (see fig. S4) and the resultant intercept of the ANCOVA in this analysis was significantly positive in the behavioural direction (value=0.219(± 0.031SE), T_(_1_9)_=7.16, p=1.24×10^-11^, BF=2.10×10^4^). Following the pre-registered hypothesis this might be interpreted as implying that the brain’s response to stimuli, requires that *at least* one oscillatory cycle having occurred in order for the bursts to be individuated.

In addition to the participant factor of the main ANCOVA, individual differences were assessed through a series of correlations between normalised behavioural performance and oscillatory amplitude applied to mean levels across frequencies *(39)* and at individual frequencies of presentation. No correlation was observed using the mean performance and amplitude, across frequencies *(40)* (r_(18)_=0.19, p=0.42, BF=0.24) or within any of the frequencies tested (see table ST1). A one sampled T-test applied to the resultant correlation coefficients, also had the capacity to demonstrate an individual difference-based relationship but did not support such a difference (T(_11_)=1.04, p=0.32, BF=0.45).

An additional post-hoc exploration of the data involved fitting 1/frequency^p^*(41, 42)* models to behavioural and oscillatory distributions. The resultant group mean β coefficient for the behavioural data was (±95% confidence bounds) 0.35(0.16) where r^2^_(11)_= 0.52 and for the oscillatory data β was 0.33(0.13) and r^2^_(_11_)_= 0.56. The tests applied indicated the β coefficients’ numerical proximity across the measures (T_(_1_9)_=1.03, p=0.32, BF=0.37). These β values correspond closely to previous reports *(60)*. The proximity of the behavioural and oscillatory β values and overlap in the confidence interval is consistent with the overall prediction and interpretation that there is a relationship between the expression of oscillatory activity and behavioural capacity to individuate information over a range of frequencies and that the distribution of the two express a similar structure. Furthermore, it was possible to probe the question of the relationship between measures in terms of individual variability via participants’ β values. This involved testing the correlation of β values, resulting from individual participants’ fits, between the two measures *(42)*, but did not support the presence of such a relationship between β coefficients: r^2^_(18)_= 0.16,p= 0.50, BF= 0.21.

**Figure S2.**
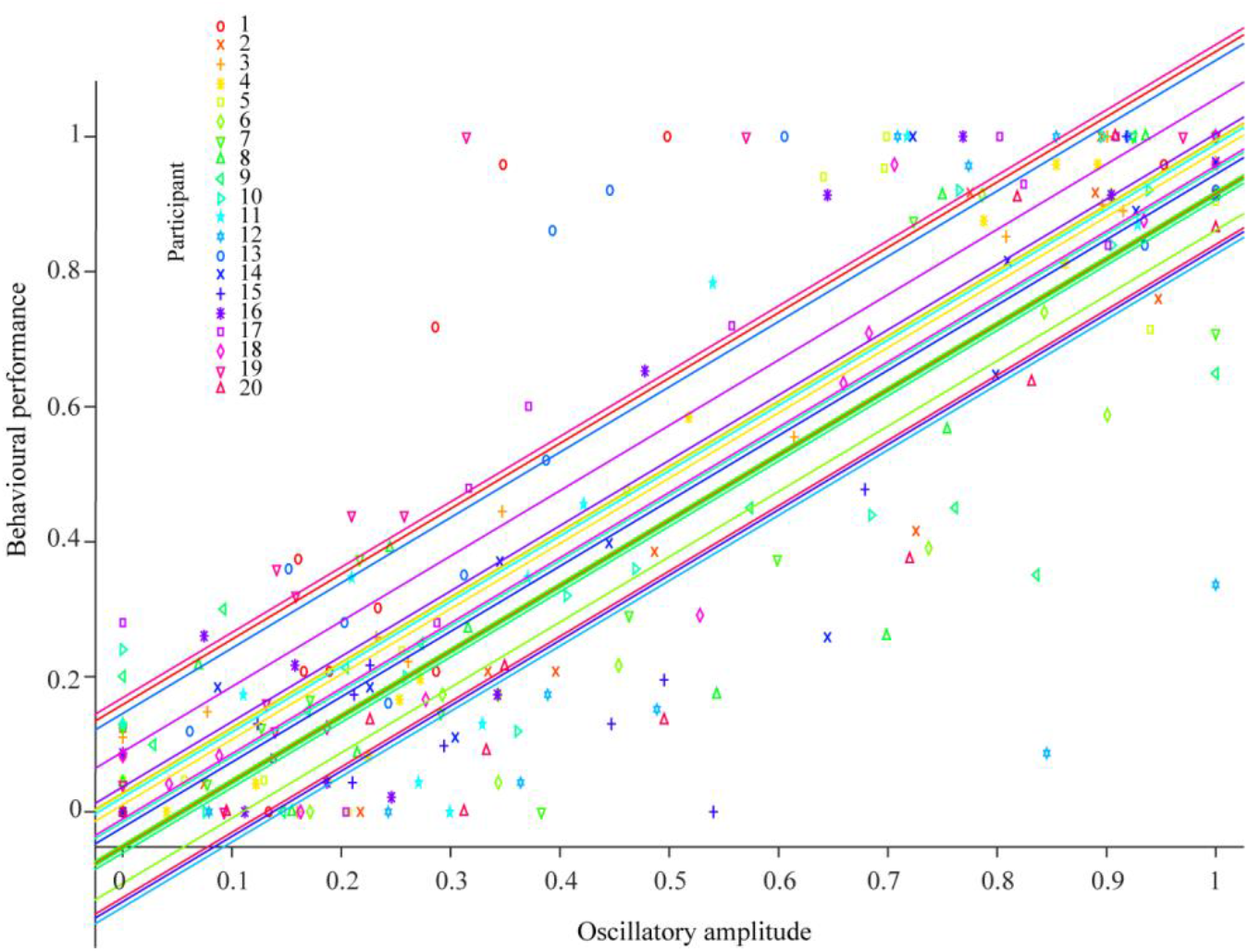
Normalised behavioural performance in burst counting against normalised oscillatory amplitude taken from the auditory cortex. Parallel lines fitted for each participant illustrating the ANCOVA *(12)*.

**Figure S3.**
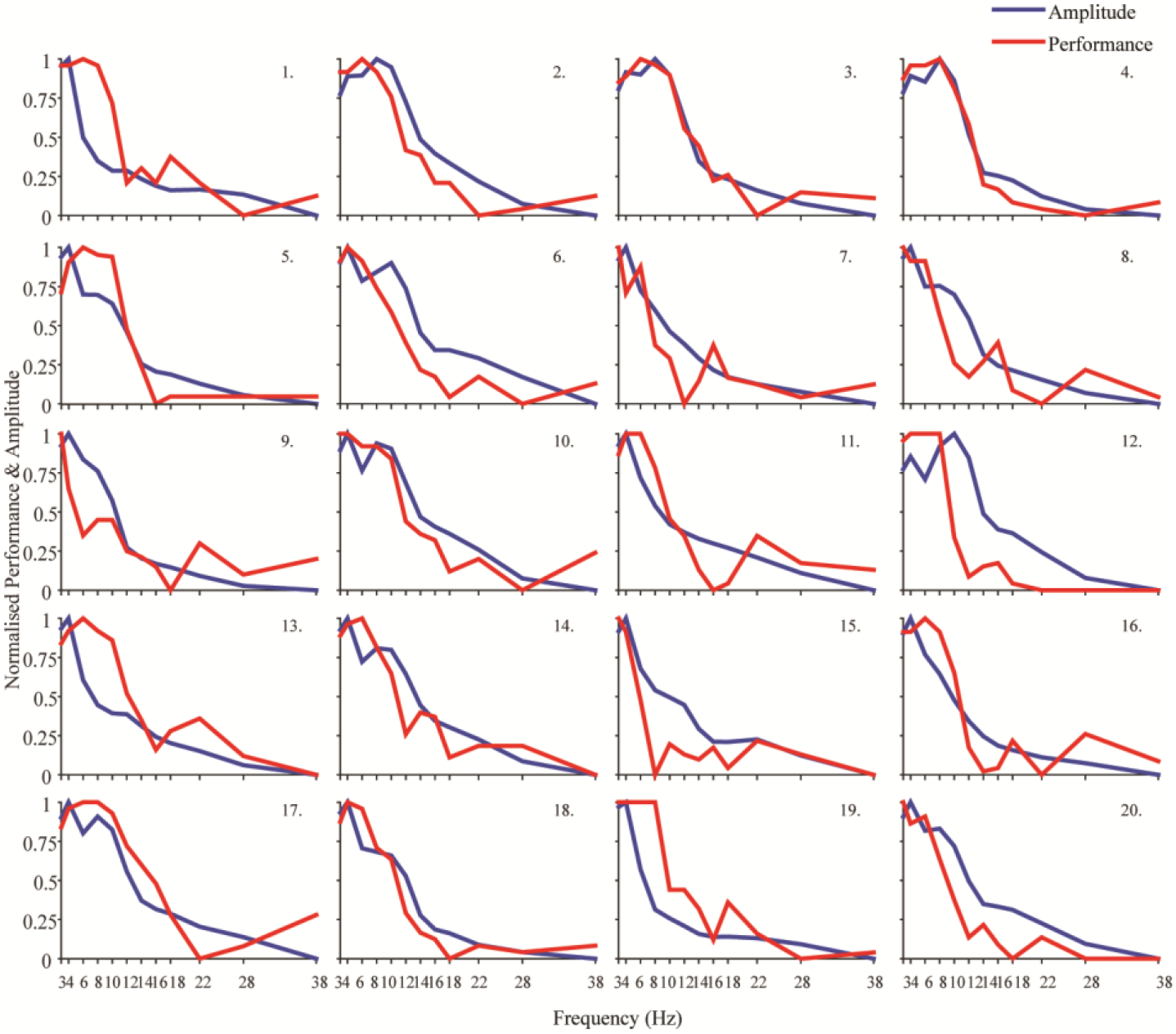
Normalised behavioural performance and normalised oscillatory amplitude for individual participants. These are the individual participants’ distributions that contributed to fig. 2.

**Figure S4.**
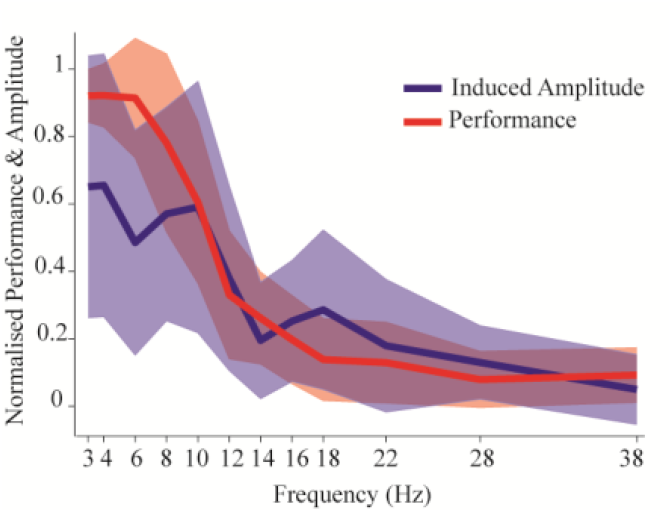
Group mean induced amplitude and behavioural performance against frequency (of presentation and oscillatory). This figure conforms to the same structure as fig. 2 of the main text but here a pre-stimulus baseline was subtracted to express the distribution of oscillatory activity in response to the stimuli.

**Table ST1.**
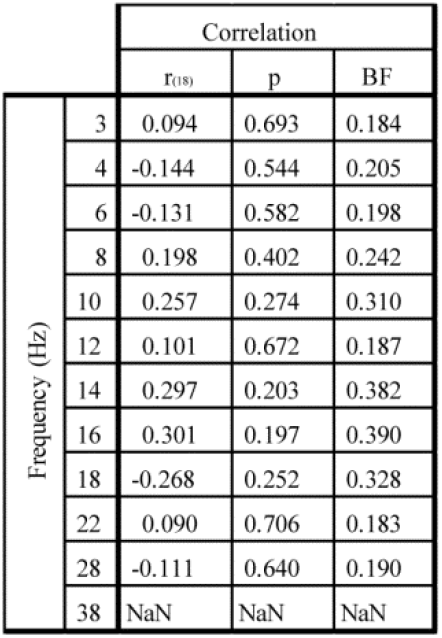
Correlation coefficient (r(df)) and corresponding Bayes Factor *(40)* for correlation between behavioural performance and oscillatory amplitude at each frequency are reported. Tests were not applicable at 38Hz as corresponding normalise oscillatory values where zero.

|  |  | Correlation |  |  |
| --- | --- | --- | --- | --- |
| | | $r_{(df)}$ | p | BF |
| Frequency (Hz) | 3 | 0.094 | 0.693 | 0.184 |
|  | 4 | -0.144 | 0.544 | 0.205 |
|  | 6 | -0.131 | 0.582 | 0.198 |
|  | 8 | 0.198 | 0.402 | 0.242 |
|  | 10 | 0.257 | 0.274 | 0.310 |
|  | 12 | 0.101 | 0.672 | 0.187 |
|  | 14 | 0.297 | 0.203 | 0.382 |
|  | 16 | 0.301 | 0.197 | 0.390 |
|  | 18 | -0.268 | 0.252 | 0.328 |
|  | 22 | 0.090 | 0.706 | 0.183 |
|  | 28 | -0.111 | 0.640 | 0.190 |
|  | 38 | NaN | NaN | NaN |

### Entrainment of oscillations by stimuli

The entrainment effect was tested with an ANOVA which compared the amplitude of the Hilbert envelope between where data was drawn at the frequency at which the stimuli were presented and compared this to the same amplitude measure but where data was drawn from other frequencies of presentation. The test of central interest here was the effect of validity reported in the main text. This relative increase in amplitude in the valid condition is within the context of a generalised (across conditions) event related desynchronisation in response to the stimuli onset *(61)*. The interaction between validity and frequency was not significant (F_(11,198)_=1.05, p=0.40, BF=0.38) which can be interpreted as the entrainment effect being present across frequencies. In line with the first set of analyses there was evidence in support of a main effect of frequency (F_(11,198)_=5.42, p=1.63×10^-7^, BF=2.02). In accordance with pre-registered criteria, the numbers of trials turned out to be insufficient to probe the potential interaction between this entrainment effect and behavioural success, so this possibility remains an intriguing avenue for future research. The BFs derived thought the supplementary application of the Bayesian ANOVA *(46)* are as follows: the main effect of validity computed as BF for main effects over the BF for the frequency main effect was 3.51. The interaction (full model over the main effects model) was 0.038 and the frequency main effect (main effects over validity main effect) was 3.90×10^6^.

As described the outcome of these analyses depended upon a randomised downsampling of trials. Therefore an exploratory analysis was undertaken to ensure that observed effects where not the result of the downsampling, involving utilisation of all available data. This supported predictions that stimuli entrained brain activity, involving relatively increased oscillatory amplitude at the frequency at which the stimuli were presented (F_(1,19)_=10.90, p=0.004, BF=11.69). Again no evidence for an interaction between validity and frequency was observed (F_(11,209)_=1.38, p=0.18, BF=0.42) and there was evidence for a main effect of frequency (F_(11,209)_=6.26 p=6.59×10^-9^, BF=2.72). The complimentary Bayesian ANOVA statistics, computed as above, are as follows: BF validity=1.88, BF frequency 2.65×10^9^, BF interaction 0.058. Figure S5 reflects the entrainment analysis applied were all trials where used as opposed to downsampled data sets reported in figure 3 of the main text.

**Figure S5.**
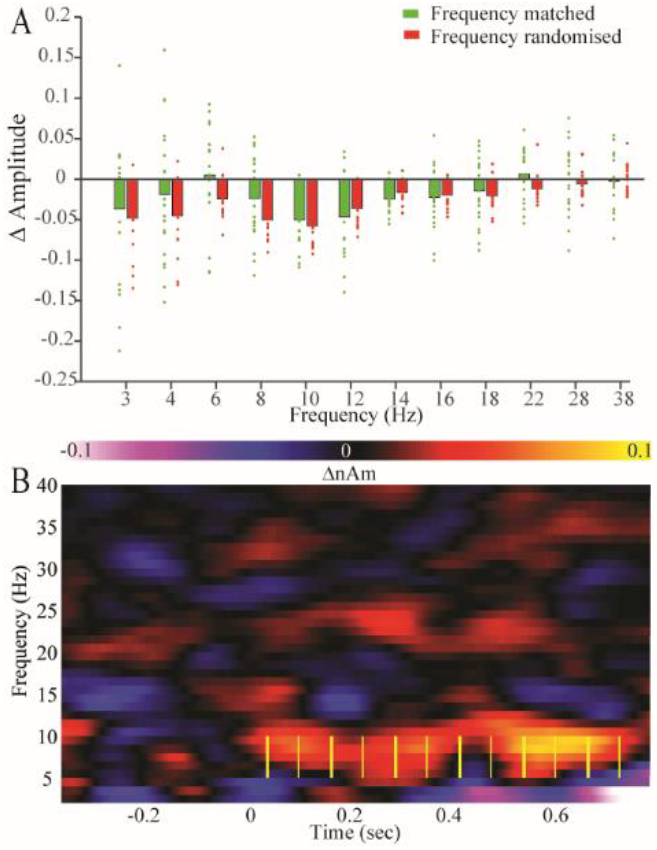
Entrainment analyses. A. Induced amplitude change, where data is drawn from either the matching frequency of presentation or all trials at other frequencies. Bars are of group mean and data points are individual participants’ data. B. Example of group level time × frequency representations of differences described in A, yellow lines illustrate the stimuli (at 8Hz here, where thicker lines represent burst onset and thinner that of silent periods). This figure conforms to the same structure as fig. 3 of the main text, but here data has not been randomly downsampled.

### Divisions of data according to behavioural success

The following analyses complement and expand upon those reported in the main text concerning successful vs. unsuccessful trials. The supplementary application of FDR correction to time × frequency analyses applied at the group level revealed three significant clusters when applied to the unbaselined amplitude difference between successful and unsuccessful trials only (see fig. S6). These differences were again within the **y** range: the first was at 37Hz and from −526 to −508ms relative to stimuli onset, the second was between 67 and 68Hz from −76 to −25ms corresponding to the time × frequency region highlighted by the mass cluster correction, and finally a 98 to 99Hz post-stimulus difference survived the FDR correction between 338 and 399ms.

**Figure S6.**
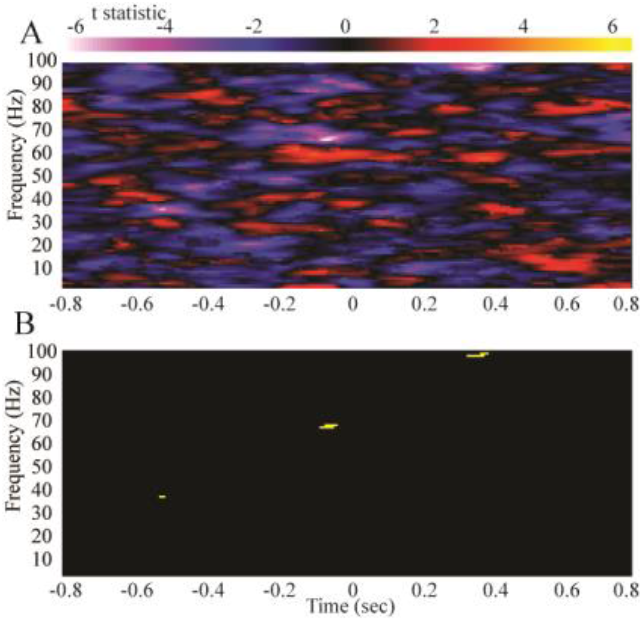
A. Time × frequency plot of t-statistic difference between successful and unsuccessful trials applied to the amplitude of the Hilbert envelope. B. Depicts the regions of these differences which surpassed false discovery rate correction for multiple comparisons *(50)*.

To test the possibility that the observed **y** differences was in part reliant upon the randomised downsampling of trials an additional set of analyses were carried out which made use of all available successful and unsuccessful trials. Although it was only possible to apply these tests to the amplitude difference using FDR correction, they indicated a similar patterning to the main registered analyses, with a significant **y** band pre-stimulus desynchronisation which survived FDR correction for multiple comparisons. As depicted in fig. S7 the FDR correction indicated that this occurred between −302 to −251ms and between 50 and 51Hz. Additionally, there appeared to be a brief post stimulus difference at 55Hz from 465 to 473ms.

**Figure S7.**
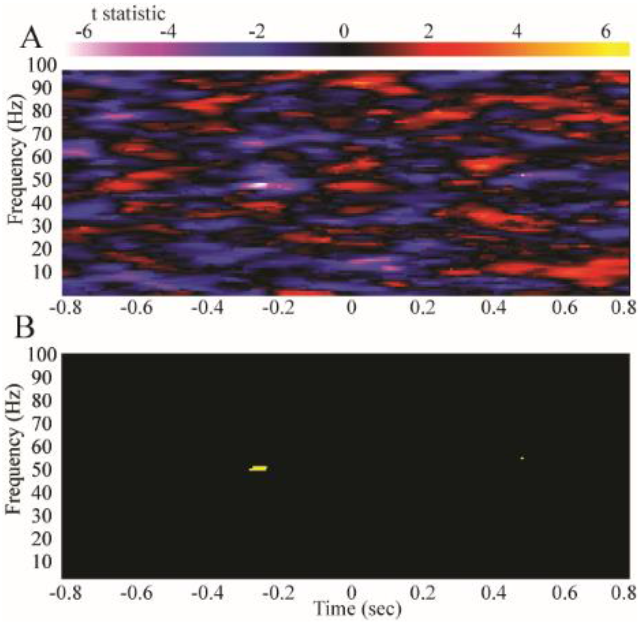
A. Time × frequency plot of t-statistic difference between successful and unsuccessful trials applied to the amplitude of the Hilbert envelope where the data to which analysis were applied consisted of all available trials rather than the downsampled balanced subset. B. Depicts the regions of these differences which surpassed false discovery rate correction for multiple comparisons *(50)*.

The oscillatory differences between when participants were and were not able to successfully separate and count the number of bursts, when explored at the group level, exposed a novel progression of **y** band changes. In addition to these main group level analyses the pre-registration document described applying comparable analyses to individual participants’ data but where the source of variance for the comparison was across trials rather than across participants. Figs. S7-S11 are the time × frequency plots of these comparisons and the summary statistics based on the mass cluster based approach are summarised in table ST2.

**S7.**
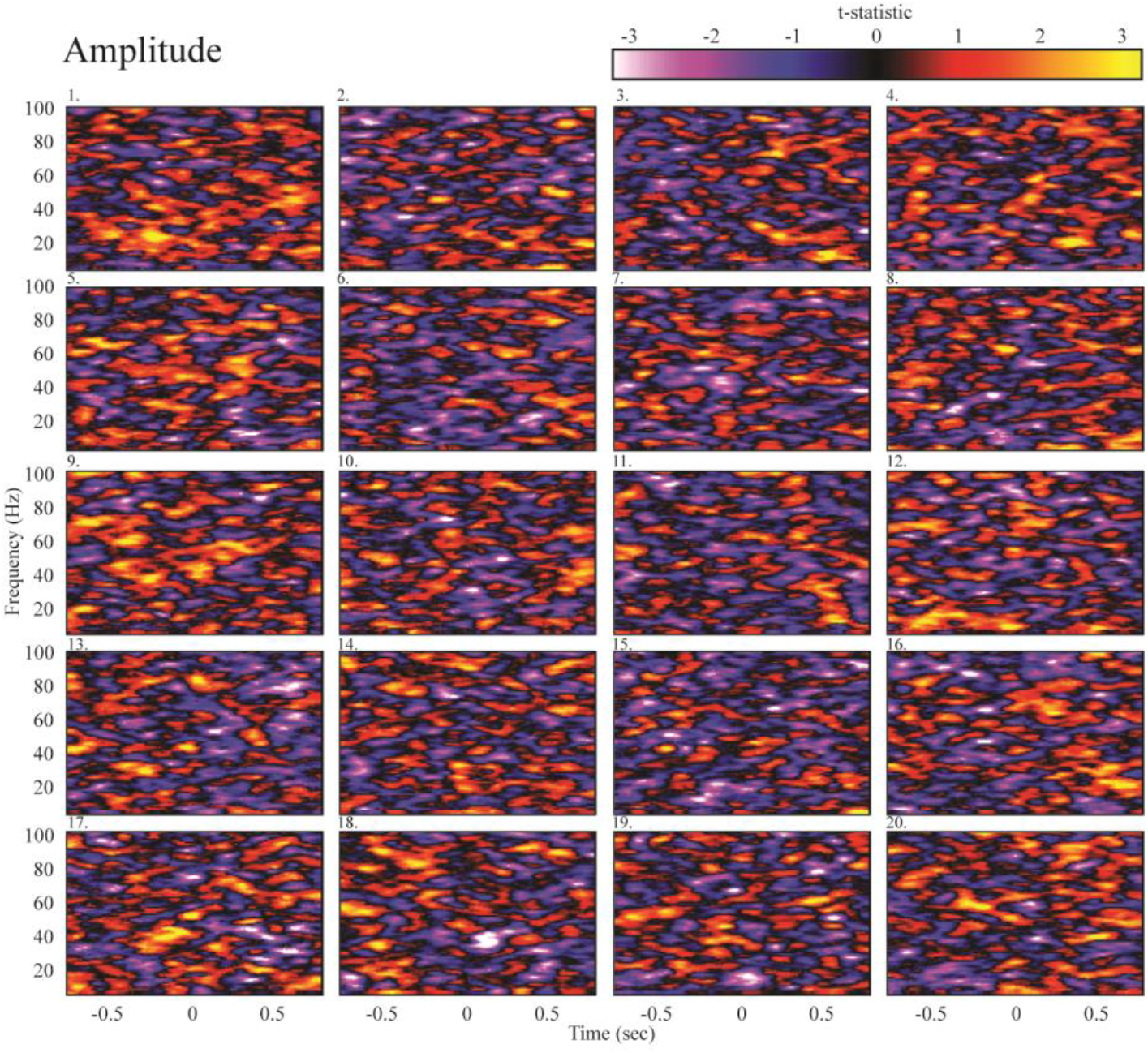
Individual participants’ time × frequency representation of amplitude difference between successful and unsuccessful burst counting performance.

**S8.**
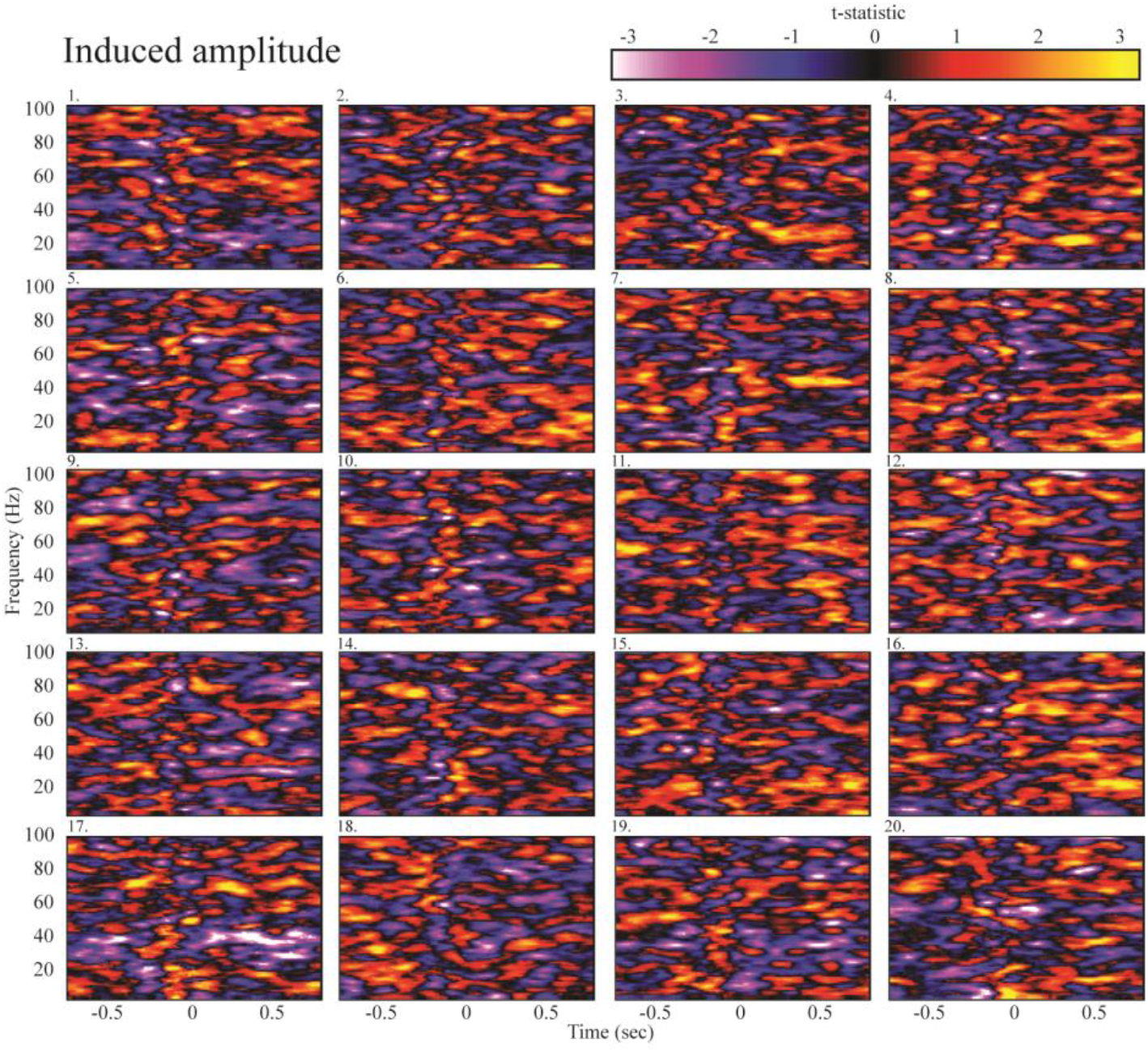
Individual participants’ time × frequency representation of induced (baselined) amplitude difference between successful and unsuccessful burst counting performance.

**S9.**
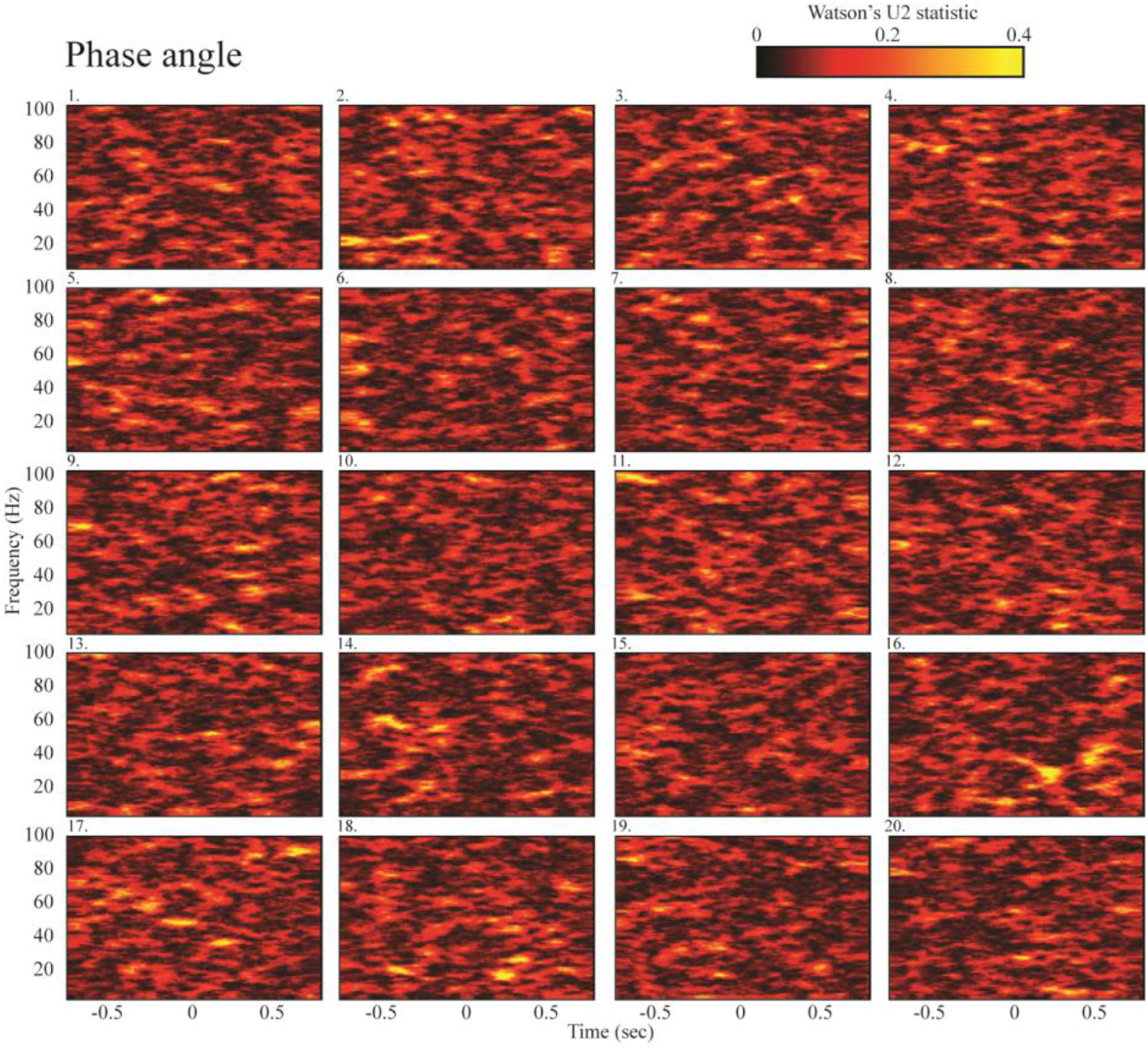
Individual participants’ time × frequency representation of phase angle difference between successful and unsuccessful burst counting performance.

**S10.**
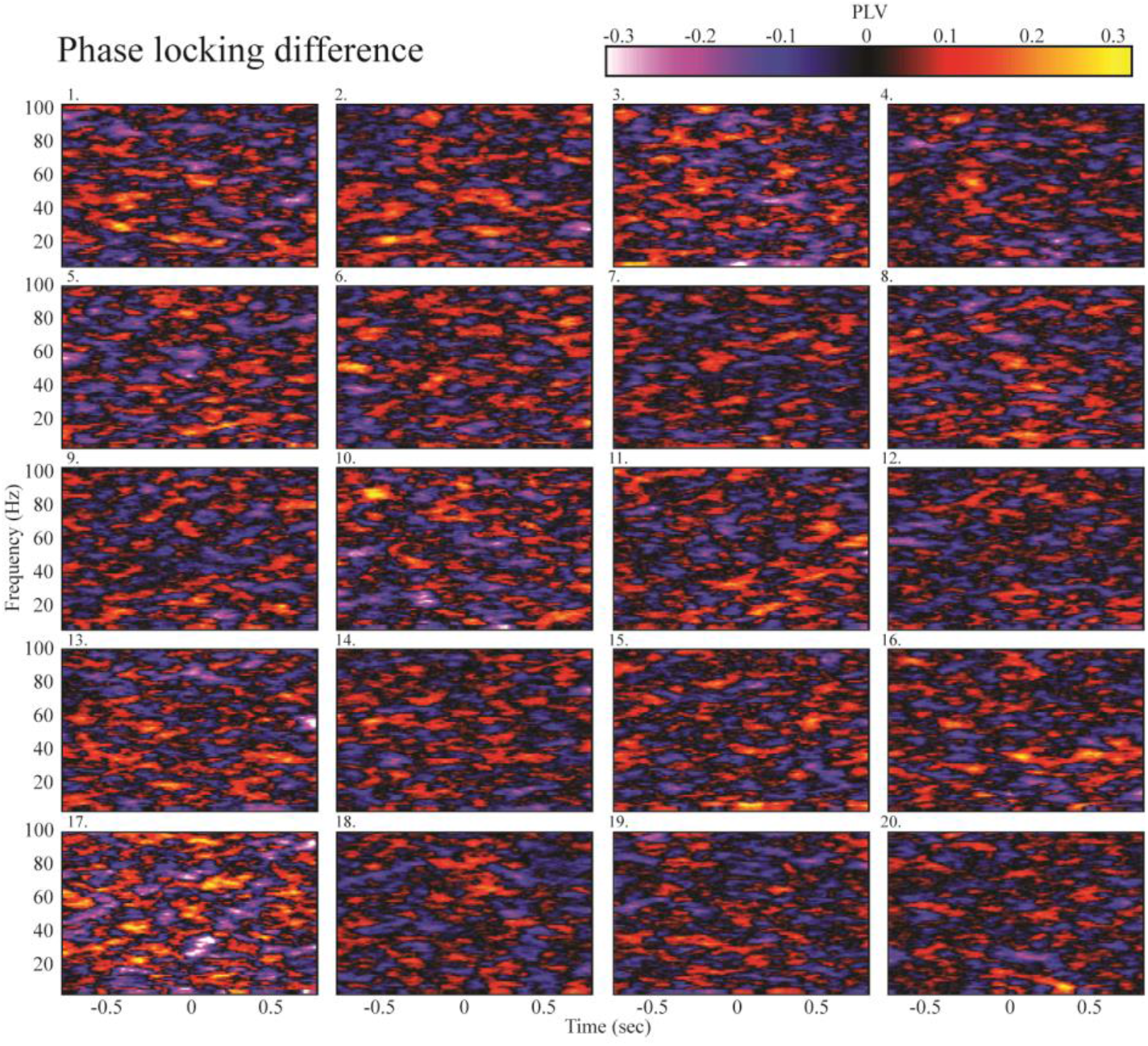
Individual participants’ time × frequency representation of phase-locking value (PLV) difference between successful and unsuccessful burst counting performance.

**S11.**
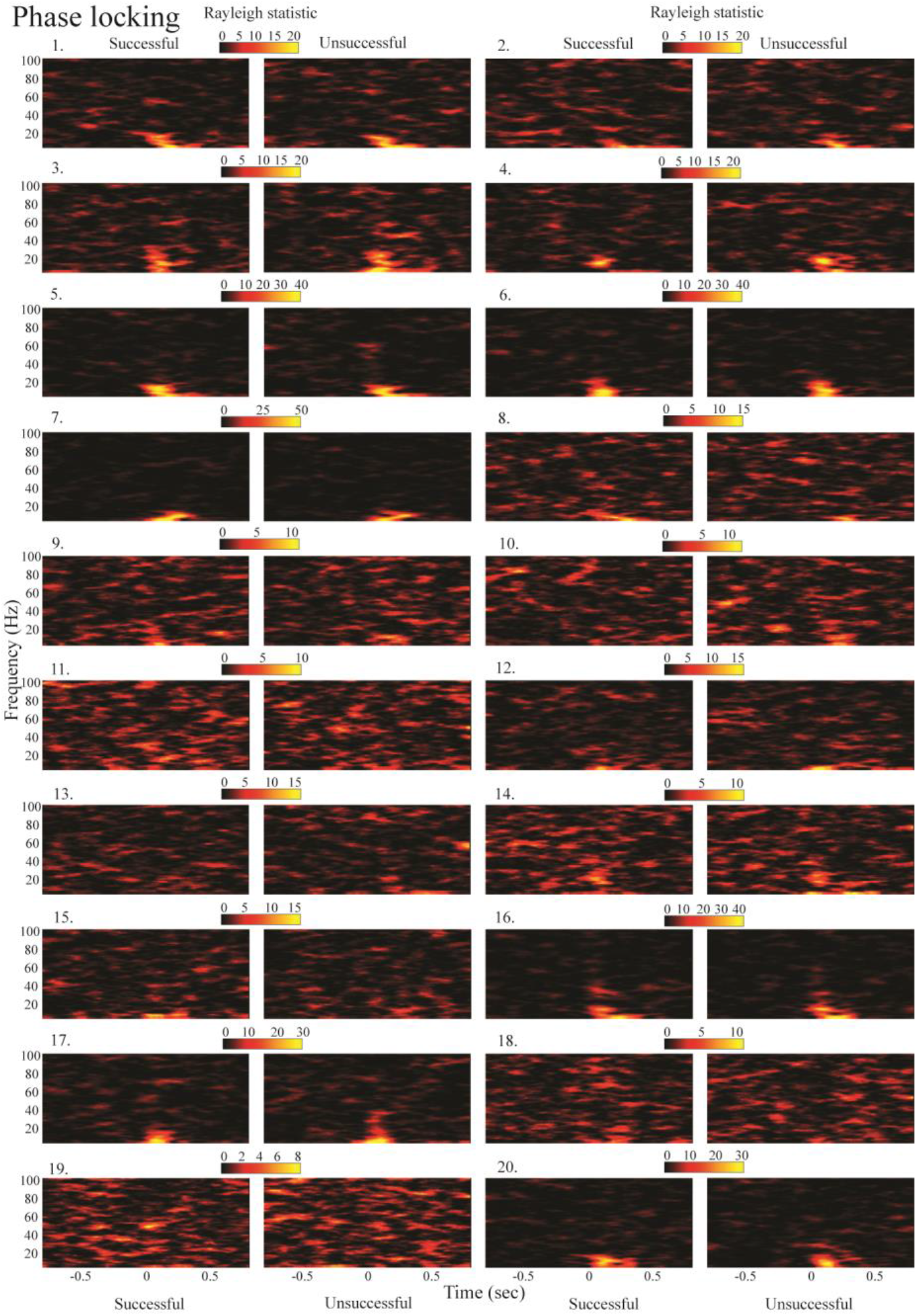
Individual participants’ time × frequency representation of phase locking in successful and unsuccessful burst counting conditions quantified via the Rayleigh statistic.

**ST2.**
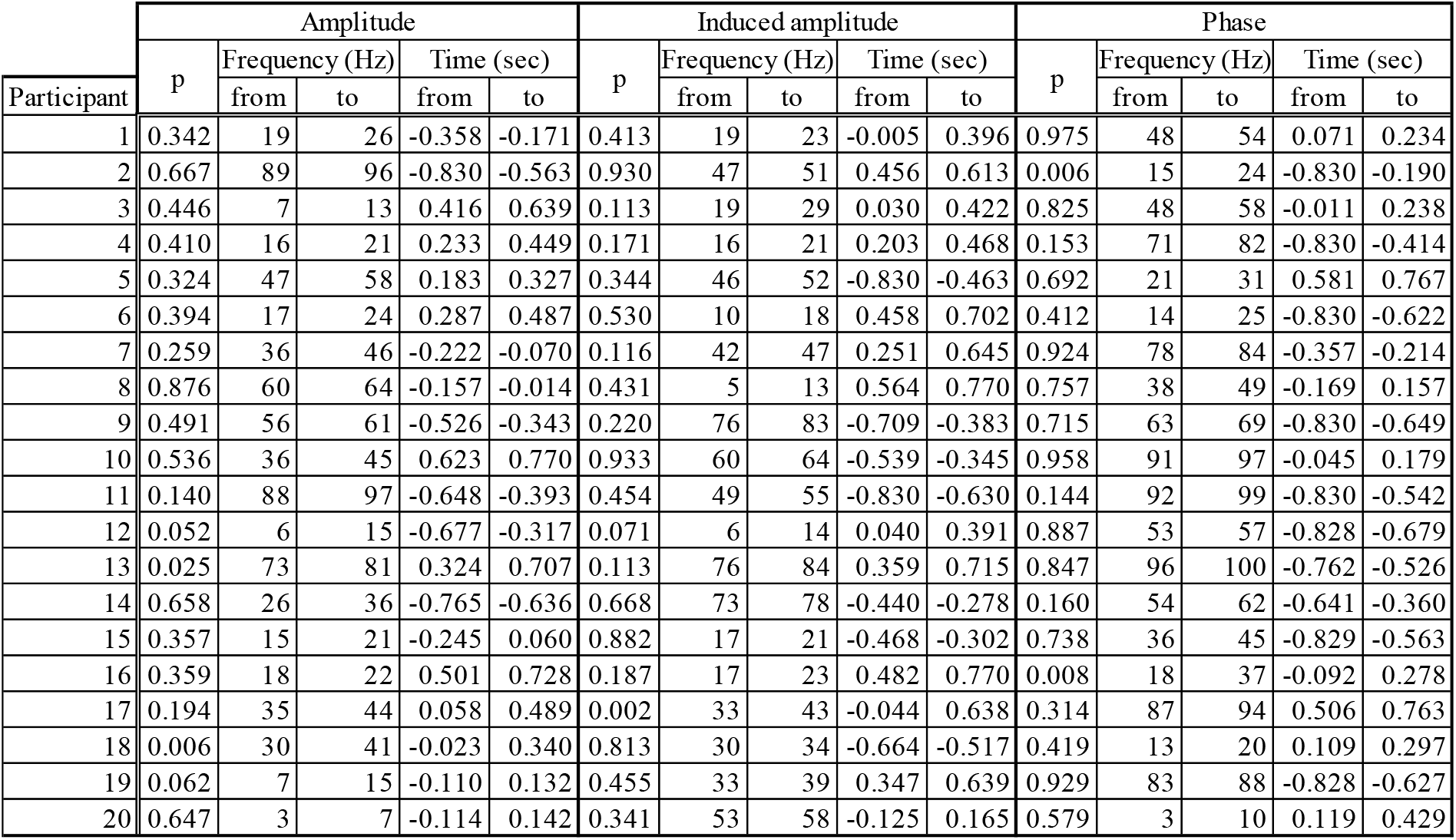
Summary of mass clusters applied at participant level. P values are the probabilities of there being a connected time × frequency region where significance criteria (p<0.05) is surpassed and has greater magnitude by chance for individual participants’ time × frequency spectra, applied to measures of amplitude difference, induced amplitude difference, phase angle. The range of the cluster (frequency and temporal range) are also provided.

The topographic distribution of the group level oscillatory amplitude differences between the successful and unsuccessful counted trials are depicted in Fig. S12. The peak location of the unbaselined pre-stimulus desynchronisation was found within the left superior temporal gyrus (Talairach coordinates: −47.2,17.1,−25.0). The largest peak of the post stimulus difference resolved to the left lentiform nucleus (Talairach coordinates: −21.1,−5.0,−7.0) just medial to the left superior temporal gyrus, where the second peak was found (Talairach coordinates: −45.2,13.1,−25.0). As the superior temporal gyrus contains the primary auditory cortex, this topography was consistent with a cortical, as opposed to muscular, origin of the observed differences. However, the peak location of the modelled dipoles resolved to a more anterior location than one might expect of pure auditory activity *(62)*. The structural-functional specificity of auditory regions has been shown to exhibit a high degree of variability *(62, 63)* and the anterior location could relate to the cognitive or semantic counting comprehension aspect of the task, activating secondary auditory areas *(64, 65)*. The proximity of the peaks to the eye does, however, still raise the possibility of an eye movement related artefact. However, the dipole appeared to be strongly left lateralised (see fig. S12). This, while consistent with a cognitive processing interpretation *(66)*, is contrary to what would be expected of an eye movement artefact, which should present bilaterally. Muscle artefacts may also be expected to extend beyond the range of the limited (<75Hz) **y** band differences observed here (see fig. 4 *56)*.

**Figure S12.**
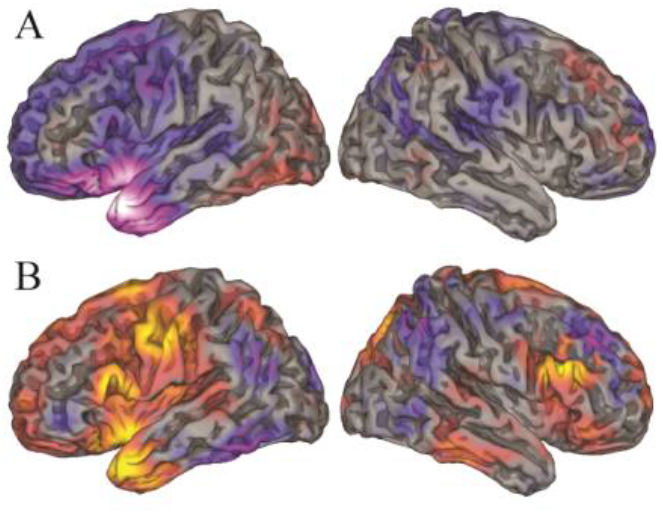
Illustrations of spatial distribution of ***y*** difference between successful and unsuccessful burst counting. Topographic representations are of SAM beamformer *(34)* solutions given parameters indicated by pre-registered analysis, for A) the unbaselined difference amplitude between performance conditions between 66-73Hz over a period from −356 to 18ms seconds relative to stimuli onset and B) Baselined (-350 to 0ms) difference between 66-73Hz over the post stimulus period 7 to 678ms. These are group means overlaid upon semi-inflated standardised brains where blue to white colours indicate relative desynchronisation in amplitude and red to yellow colours indicate relative increase in synchrony (see mri3Dx (Singh KD)).

As clarification with respect to the interpretation of the **y** band observations: The three measures which expressed sequential events in the **y** range suggest such oscillations may constitute, contribute or signify a particular state which is conducive to the individuation of content and task performance, as described in the main text. Previously it has been suggested that certain phases of oscillatory cycles, relative to the onset of stimuli, might allow information to be passed *(22, 4, 18)*. However, these previous demonstrations have tended to concern oscillatory activity around the **a** range, where a relatively low number of oscillatory cycles might encompass the time × frequency region expressing a difference in capacity to individuate content. The interpretation often applied is that a certain part of the oscillatory cycle is conducive to performance (e.g. *22)*. Here, by contrast, the relatively fast **y** band cycles mean multiple oscillatory cycles are completed, covering many phases, over the region where the phase contrast expressed differences between the successful and unsuccessful conditions. While this does not rule out the possibility that a certain phase, at a certain time, may be conducive to content individuation, neither does it directly suggest it. Rather the suggestion from the current data is that there is a state difference, involving phase angle, between the conditions, that is conducive to individuation of bursts, but precisely what the conducive phases are is not clear from the present data. The additional possibility of the **y** progression observed being coupled to particular phases in low frequencies (e.g. *18, 67)* is an area for further investigation.

There are three additional noteworthy points with respect to the progression of the **y** band events. First the induced post-stimulus synchronisation when pre-stimulus baseline is subtracted may, at least in part, be due to the subtraction of the relative desynchronisation, apparent in the raw amplitude contrast, prior to the onset of the stimuli. Second, although no differences in the phase consistency contrast were apparent, when corrections for multiple comparisons were applied, there did appear to be a possible spike fluctuation in phase constancy in the successful condition at around 70Hz and at approximately 300ms prior to stimulus onset (see(fig. 4D). Normally such a fluctuation would be not merit note. However, because it appears at the same frequency to that of other demonstrated effects and, intriguingly, at the boundary between the phase and amplitude difference, its potential occurrence highlights it as a possible target for future research. Finally, within recent theories of predictive coding, a link has been made between **y** band responses, as prediction error, to feed-forward/bottom-up processing, i.e. in responses to stimuli *(68, 69)*. However, as the **y** progression observed here starts prior to the onset of the stimuli, there would appear to be an inconsistency. Although there are ameliorating explanations, the basic incongruity is noteworthy in a context of predictive coding where testable unique predictions are scarce *(70)*.

### Participant questionnaires

Participants responded to questionnaires following their participation in the experiment. These were intended to probe individual differences in performance strategy, which could then potentially have been related to differences in the MEG data. The questions are given at the end of the methods section and responses given are provided in table ST3.

To summarise the responses collected using the questionnaires: All participants responded that they “counted the numbers of bursts in their heads” (Q1A) and did so on the majority of trials (Q1B: mean proportion (±standard deviation (SD)): 0.81±0.152). A greater number of participants reported that they counted the stimuli as they were presented (12/20) compared to referring back after presentation (3/20), although 5 participants reported using both strategies (Q2A). When asked about how often they adopted the strategy of reflecting back afterward, the responses were highly variable (Q2B: 0.43(±0.20SD)). Use of the duration of the burst trains as a strategy was reportedly adopted by the majority of participants (Q3A: 15/20), but only on a relatively low proportion of trials (Q3B: 0.24(±0.22SD)) and participants reported almost exclusively doing so when stimuli were presented at the higher frequencies (Q3C: 15/16).

Table ST3 summarises the correlations applied between the experimental measures and responses participants gave to the questionnaires. Two correlations passed the frequentist criteria level of p<0.05 without correction for multiple comparisons. One of these was a potential relationship which involved a negative correlation between the magnitude of the induced **y** difference between successful and unsuccessful trials and the extent to which participants adopted the strategy of estimating the number of bursts based on the duration of burst trains (question 3B). However, this would not survive correction for multiple comparisons and did not demonstrate substantial evidence in favour of an association via the Bayesian method. Therefore no conclusions should be drawn, although it may highlight a potential area for future investigation. The other apparent correlation would not survive Bonferroni correction for multiple comparisons either, but the corresponding BF (8.18) did indicate substantial evidence in favour of an association was between the extent to which participants ‘counted the number of bursts in their heads’ (1B) and the gradient (or association) between the behavioural and oscillatory primary measures. This relationship was predicted and potentially suggests that participants who adopted the ‘counting burst’ strategy may have expressed a more pronounced correspondence between behavioural and oscillatory measures, which in turn is consistent with the overall interpretation of the experiment where individuation is supported by oscillations. However, the analyses of this section are all exploratory and as such should be treated with caution. Furthermore it is apparent that the questionnaires may have failed to capture important differences of interest i.e. the phenomenology of the task. Therefore a potential avenue for further investigation with respect to tasks such as this, would be a thorough investigation of the structure of the experience of the task, linking fine grain differences in experience to MEG measures *(71, 72)*.

---

**Table ST3.** Summary of participants’ responses to questionnaires. ‘Nan’ indicates no response was entered.

| Participant | Questions |  |  |  |  |  |  |  |  |
| --- | --- | --- | --- | --- | --- | --- | --- | --- | --- |
|  | 1A | 1B | 2A | 2B | 3A | 3B | 3C | 4 | 5 |
| 1 | 1 | 0.8 | 2 | nan | 1 | 0.05 | 2 | nan |  |
| 2 | 1 | 0.95 | 2 | nan | 1 | 0.3 | 2 | nan | Visualized bursts as a jagged line. |
| 3 | 1 | 0.9 | 2 | 0.2 | 1 | 0.2 | 2 | 0.25 |  |
| 4 | 1 | 0.8 | 3 | 0.4 | 1 | 0.1 | 2 | 0.1 | Often unsure between 6 or 7, at times would guess 7 only because I had not done that response often. |
| 5 | 1 | 0.9 | 2 | 0.65 | 1 | 0.4 | 2 | 0.2 |  |
| 6 | 1 | 0.8 | 2 | 0.1 | 1 | 0.1 | 2 | 0 |  |
| 7 | 1 | 0.4 | 1 | 0.75 | 1 | 0.8 | 2 | 0 |  |
| 8 | 1 | 0.5 | 2 | nan | 0 | nan | nan | 0.2 |  |
| 9 | 1 | 0.85 | 1 | 0.55 | 1 | 0.4 | 2 | 0.15 | Repeat sound in my head. Kept thinking I was missing a number when counting -usually 1 or 2. |
| 10 | 1 | 0.7 | 2 | nan | 0 | nan | nan | 1 | None - just concentrate as much as I could |
| 11 | 1 | 0.9 | 2 | 0.75 | 1 | 0.4 | 2 | 0.1 |  |
| 12 | 1 | 1 | 3 | 0.4 | 0 | nan | nan | 0.1 | Compared current burst to previous one and noted if it was shorter or longer (only if similar frequency) |
| 13 | 1 | 1 | 2 | 0.5 | 0 | 0 | 3 | nan |  |
| 14 | 1 | 0.8 | 1 | 0.5 | 1 | 0.25 | 2 | 0.25 |  |
| 15 | 1 | 0.7 | 3 | 0.5 | 1 | 0.5 | 2 | 0 |  |
| 16 | 1 | 0.9 | 3 | 0.4 | 1 | 0.15 | 2 | 0.05 |  |
| 17 | 1 | 0.8 | 2 | nan | 1 | 0.05 | 2 | 0.1 |  |
| 18 | 1 | 0.95 | 2 | 0.2 | 1 | 0.1 | 2 | 0.1 |  |
| 19 | 1 | 0.8 | 3 | 0.5 | 1 | 0.15 | 2 | 0.1 | When it was slow, I counted directly, if too fast I had to reflect back in my head. |
| 20 | 1 | 0.9 | 2 | 0.1 | 0 | 0 | nan | 0 | Counted as in music 1 and 2 and etc |

**Table ST4.**
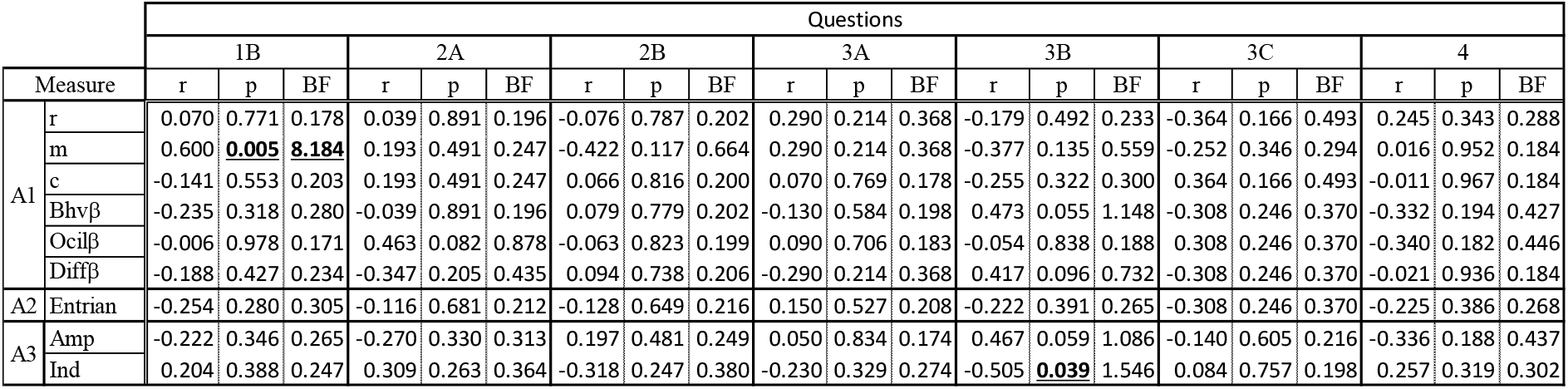
Summary of correlations applied between experiment measures and questionnaire responses. r,m and c are the results of the correlations between the behavioural and oscillatory distributions described in the first analysis. **p** refers to the **P** coefficients applied in exploration of the oscillatory (Ocil) and behavioural (Bhv) distributions and ‘Diff refers to the difference between these **p** values. Entrain refers to the quantification of the entrainment of oscillatory responses by the stimulus, investigated in the second main analysis and the final set of measures represent the differences in amplitude between the successful and unsuccessful trials in the induced (Ind) response to stimuli and without a trial level baseline (Amp). Questions 1B-4 are described above. Correlation r,p and BF are provided for each comparison. Highlighted are correlations where there is evidence for some form of relationship.

## Acknowledgments

This work was completed with the aid of grants from the Biotechnology and Biological Sciences Research Council (grant code BB/K008277/1) and the Wellcome Trust (grant code 104943/Z/14/Z). I would also like to thanks the following people for their help with this project: Chris Chambers, Ralph Connor, Emily Hammond, Suresh Muthukumaraswamy, Krish Singh, Catherine Walsh.

